# Hyaluronan Underlies the Emergence of Form, Fate, and Function in Human Cardioids

**DOI:** 10.64898/2026.01.26.700345

**Authors:** Stefan M. Jahnel, Anna Dimitriadi, Julia Kodnar, Vasileios Gerakopoulos, Yajushi Khurana, Maximilian Mayrhauser, Tobias Ilmer, Mohamed Amin Aguech, Adam R. Hall, Keisuke Ishihara, Sasha Mendjan

## Abstract

The extracellular matrix (ECM) is crucial for organ development and disease. Yet, the interplay among cells, function, and the ECM during human cardiogenesis remains obscure. Using human cardioids, we discovered that cardiac mesoderm-synthesized hyaluronan (HA) underlies early cardiac functional development. HA drives cardioid cavity formation through hydrogel swelling and bioscaffolding, critical functions of the cardiac jelly in the early vertebrate heart. During an early developmental window, HA is essential for establishing cardiac cell identity, while at later stages, HA-generated forces promote beating function through mechanosensitive channels. Chamber-specific differences in mechanical sensitivity ensure robust contractions in multi-chambered tissues. Our findings reveal how a single endogenous ECM component orchestrates the co-emergence of form, fate, and function during human organogenesis, opening new avenues for bioengineering physiologically relevant organ models.

## INTRODUCTION

Organogenesis unfolds through dynamic interactions between cells and their physicochemical environment (*1*). Progenitors produce the extracellular matrix (ECM), which, in turn, provides chemical and mechanical cues that direct cellular behavior (*2–4*). This cell-ECM feedback is a fundamental principle that underlies the emergence of functional organs. The first organ to form and function in the human embryo is the heart (*5–8*), which starts to beat around embryonic day 20 (*9*). The ECM is crucial for the functional development of the heart (*10*, *11*), providing both biochemical signals that regulate the behavior of diverse cardiac cell types and the mechanical scaffolding that supports the heart’s rhythmic contraction. While numerous ECM components have been identified in cardiac development and disease (*12*, *13*), the precise functional dissection of cell-ECM interactions orchestrating human cardiogenesis remains challenging.

The inaccessibility of the developing human embryo has long prevented direct mechanistic studies of human cardiogenesis. Recent advances in three-dimensional models of the human heart derived from human pluripotent stem cells (hPSCs) provide a powerful experimental platform, recapitulating self-organized chamber formation, regional identity specification, and contractile function *in vitro* (*14–17*). However, these systems have not been used to study the role of endogenous cardiac ECM, despite its central role in development and disease. Here, we leverage chamber-like human cardiac organoids (*14*, *15*), or cardioids, to dissect how form, fate, and function emerge during early heart development. We discover that cell-synthesized hyaluronan (hyaluronic acid, HA) drives chamber morphogenesis, promotes cardiomyocyte differentiation, and establishes a mechanical microenvironment that supports the earliest heartbeats. Our findings reveal how HA, a key ECM component, coordinates the co-emergence of structure, identity, and function during human organ development.

## RESULTS

### Cardioids generate cavities that contain hyaluronan

Fluid-filled cavities are the hallmark of the chambered architecture of cardioids. While our previous studies showed cavities emerge by d2.5 (*14*, *15*) (Fig. 1A), bright field microscopy (Fig. 1B) and cryo-sections provide limited understanding of 3D tissue structure. Therefore, we focused on left ventricle (LV) cardioids that generate the largest cavities of all cardioids (*15*) and performed whole-mount 3D imaging and volumetric analysis (Methods, Fig. 1C). We found that cardioids undergo a pancake-to-spherical shape transition from d1.5 and the subsequent 12 hours. From d2.5 onwards, cardioids adopt spherical morphology with multiple cavities. The total volume of cardioids expands rapidly between d2.0 and d4.5, subsequently plateauing at d5.5 (Fig. 1C’), with cavities comprising 30-40% of total volume between d2.5 and d5.5 (fig. S1A). The number of cavities increased over time (Fig. 1C’). To monitor the dynamics of cavity formation, we performed live volumetric imaging on CAG-GFP cardioids and observed that multiple cavities emerge, grow, and coalesce within cardioids (Fig. 1D, 1D’, fig. S1B, Movie S1, S2, S3). Scanning electron microscopy revealed a loosely connected cell arrangement (Fig. 1E), suggesting that the tissue is porous. To test the permeability of molecules in cardioids, we added fluorescently labelled Dextran to the culture media. Within 10 min, 150 kDa Dextran rapidly permeated the cavities (Fig. 1F). Thus, unlike mammalian blastocysts (*18*, *19*) or many other organoids (*20*, *21*), cardioid cavities rapidly expand in the absence of a tightly sealed tissue.

**Figure 1.**
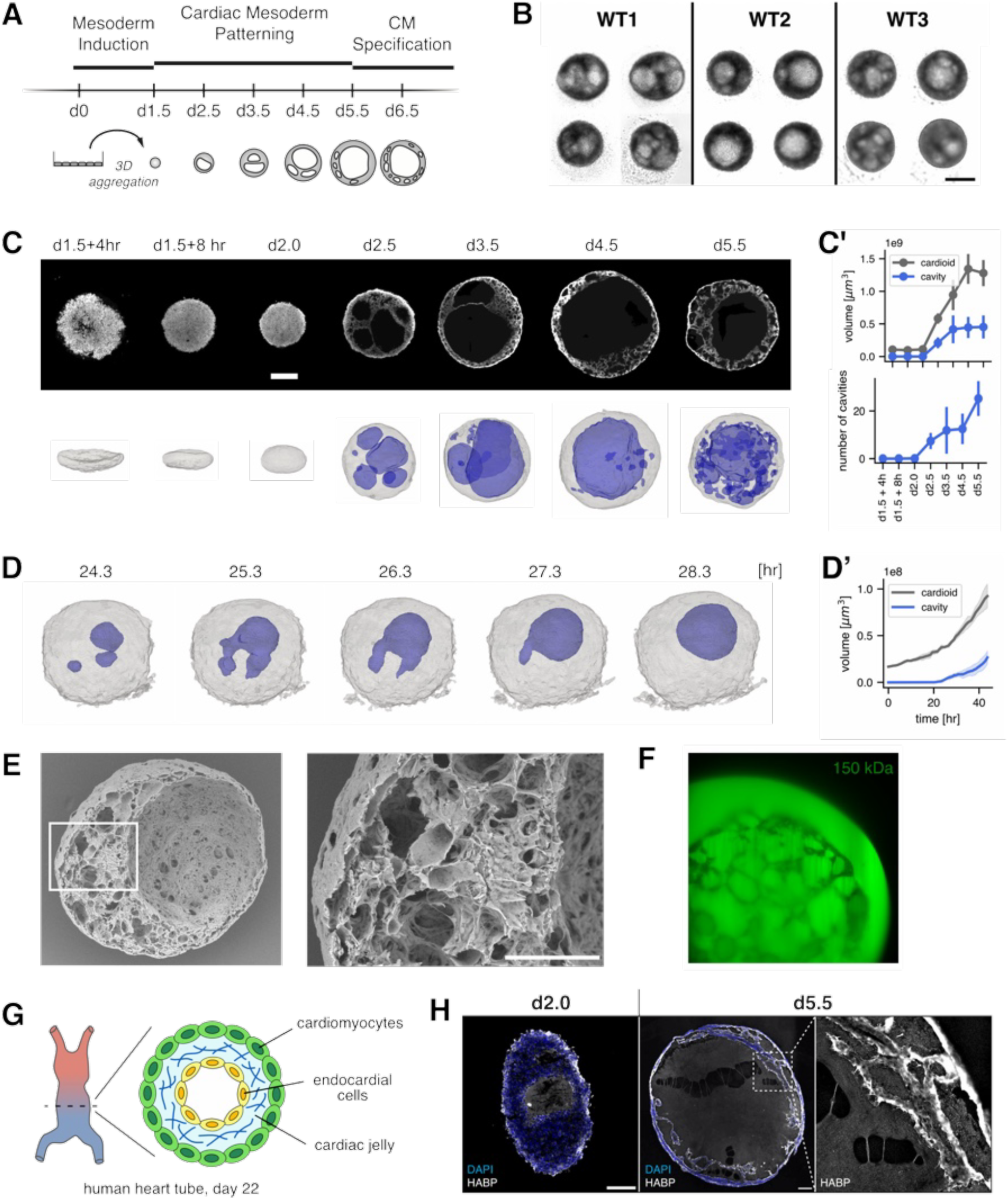
Cardioids generate cavities that contain hyaluronic acid. (A) Schematic of cardioid development and time course of cavity formation. (B) Brightfield images of left ventricular cardioids at day 2.5 containing cavities. Examples are shown for cardioids generated from three cell lines. (C) Top: Confocal images of cardioids labeled with AlexaFluor 647. Bottom: Surface renderings of the outer cardioid boundary (gray) and cavities (blue) of the same samples. Scale bar 400 micron. (C’) Top: Volume of cardioid and cavity at each time point. Bottom: Number of cavities per cardioid at each time point. Error bars indicate the standard deviation across n=4-12 samples per time point. (D) Surface renderings of selected time points from the live microscopy data (Movie S2, S3), where 0 hr was defined as the beginning of the imaging. (D’) Volume quantification of cardioid and cavity of the live microscopy date. Error bars indicate the standard deviation across n=4 samples. (E) Scanning electron microscope images show the porous nature of cardioids. Scale bar 100 micron. (F) Confocal image of a cardioid (d4.5) incubated with 150 kDa FITC-Dextran overnight. (G) Schematic of the human heart tube at day 22. Similar to the heart tube of other vertebrates, the cardiac jelly fills the space between the myocardium and endocardium. (H) Cryo-section images of cardioids showing the localization of hyaluronic acid binding protein (HABP). Scale bar 100 microns.

How can such a porous tissue increase the volume in its acellular space? We considered the developmental mechanism employed by the vertebrate heart tube, where the large extracellular space between the endocardium and myocardium is occupied by a gelatinous substance termed the ‘cardiac jelly’ (*22*, *23*), which largely consists of hyaluronic acid (HA) produced by cardiac mesoderm and cardiomyocyte progenitors (*24–26*) (Fig. 1G). The swelling of the HA hydrogel leads to the rapid volumetric expansion of the heart tube. To investigate whether a similar mechanism may operate in cardioids, we performed histological staining and found that HA is present throughout the cavities (Fig. 1H). To independently confirm HA identity and characterize its molecular weight, we performed single-molecule solid-state nanopore analysis on HA extracted from day 6.5 LV cardioids, revealing a mean molecular weight of 1,243 kDa (fig. S1C).

### Hyaluronan synthesis by HAS2 drives cavity formation in cardioids

To test if HA plays a role in shaping cardioids, we enzymatically degraded HA using hyaluronidase (HAase). When d2.5 left ventricular cardioids were treated with HAase, they began to shrink (Fig. 2A), with diameters decreasing by 22 % after 10 hours (Fig. 2A’). Sections of cardioids revealed complete collapse of cardioid cavities following HAase treatment. (Fig. 2B, Movie S4). HAase treatment at d3.5, 4.5, and 5.5 showed a reduced yet consistent decrease in cardioid size (Fig. 2A’, Movie S5). These results suggest HA exerts an outward force that counteracts cell contractility to sustain cavity volume.

**Figure 2.**
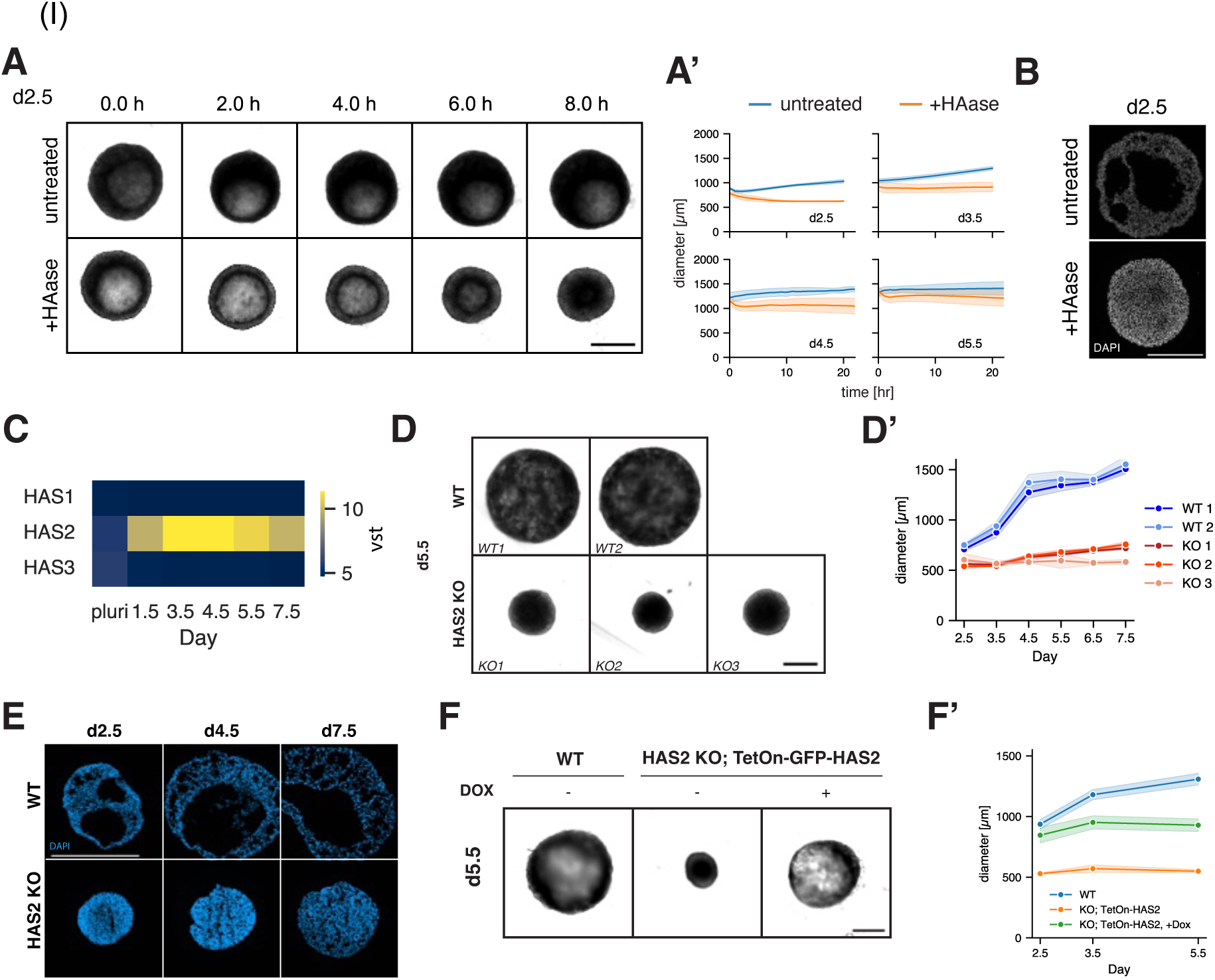
Hyaluronic acid synthesis by HAS2 drives cavity formation in cardioids. (A) Brightfield image sequences show that hyaluronidase (HAase) treatment results in rapid shrinkage of day 2.5 cardioids and their cavities, while untreated cardioids continue growth. Scale bar 500 microns (A’) Size reduction with HAase treatment is quantified for cardioids at day 2.5 to 5.5. n=8 samples per condition. Shaded regions indicate standard deviation. (B) Cryo-section images of untreated and HAase-treated (treatment d1.5-2.5) day 2.5 cardioids. Scale bar 250 microns. (C) Heatmap visualization of the transcripts detected for HAS1, HAS2, and HAS3 genes during cardioid development. VST, variant stabilized transcript counts. (D) Brightfield images of wild type (WT) and HAS2 KO cardioids at d5.5. Scale bar 500 microns. (D’) Diameter of cardioids derived from WT and HAS2 KO genotypes. Colors distinguish samples prepared from n=2 WT cell lines and n=3 HAS KO cell lines. n=15-16 samples per cell line per timepoint. Shaded regions indicate standard deviation. (E) Representative cryo-section images of WT and HAS2 KO cardioids showing absence of cavities in HAS2 KO. Scale bar 500 microns. (F) Brightfield images of WT and HAS2 KO; TetOn-GFP-HAS2 cardioids at day 5.5. Doxycycline was supplemented to the medium from day 1.5 to 5.5 for the indicated condition. Scale bar 500 microns. (F’) Diameter of cardioids derived from WT and HAS2 KO; TetOn-GFP-HAS2 genotypes. n=11-12 samples per cell line per timepoint for day 2.5 and 3.5. n=5 samples per cell line per timepoint for day 5.5. Shaded regions indicate standard deviation.

While the human genome encodes three members of the hyaluronic acid synthase family of enzymes, HAS2 is the only known isoform implicated in vertebrate heart development (*25–27*). Through transcriptomic profiling, we determined that HAS2 is the only synthase that is expressed during cardioid differentiation (Fig. 2C). To test if HAS2 is responsible for HA production, we established three independent HAS2 KO lines and generated cardioids (fig. S2A, B). HAS2 KO cardioids remained small in size relative to their WT counterparts throughout differentiation and did not exhibit readily observable cavities in brightfield images (Fig. 2D, Fig. 2D’, fig. S2C). Histological sections confirmed the absence of cavities in HAS2 KO cardioids (Fig. 2E). To test whether the morphology of HAS2 KO cardioids can be rescued genetically, we engineered a chemically inducible system for GFP-HAS2 expression in the HAS2 KO background (HAS2KO; TetOn-GFP-HAS2). Upon addition of doxycycline from d1.5, we observed GFP-HAS2 expression (fig. S2D) and an increase in cardioid size from d2.5 to d5.5 (Fig. 2F, Fig. 2F’). In summary, we identified a HA hydrogel, produced by HAS2, as the main driver of cavity formation in cardioids.

### Hyaluronan is necessary for cardiac mesoderm specification

Animal studies have shown that HAS2 is essential for normal heart development and embryonic viability (*25*). Cardioid development is characterized by a transition from nascent mesoderm through cardiac mesoderm into functional cardiomyocytes (*14*) (Fig. 3A). To investigate the role of HA during LV cardioid development, we harvested HAS2-KO cardioids at different timepoints (d1.5, d2.5, d4.5, d7.5) and performed bulk RNA sequencing. Up to d1.5 HAS2-KO had no effect on the transcriptome. From d2.5 onward, the transcriptome of HAS2 KO samples diverged sharply from that of WT controls (Fig. 3B). To test if enzymatic degradation of HA has a similar effect to genetic deletion of HAS2, we started HAase treatment from d1.5 and followed with transcriptomic profiling (Fig. 3C). At the transcriptomic level, HAase-treated cardioids largely phenocopied stage-matched HAS2 KO cardioids at d2.5 (Fig. 3D). Both KO and HAase-treated cardioids downregulated pathways associated with cardiac muscle development and function attributed to a misregulation of typical cardiac regulatory (e.g. TBX5, NKX2-5, GATA4) and structural (e.g. TNNT2, TPM1, ACTC1, MYL7, MYL4, MYH9) genes (Fig. 3D, Fig. 3E, fig. S3A, fig. S3B, fig. S3C, Table S1, Table S2). On the other hand, KO and HAase-treated cardioids showed prolonged expression of genes linked to early mesoderm development (e.g. MESP1, MESP2, MIXL1, CER1) and a bias towards a paraxial/intermediate mesodermal (e.g. TCF15, DLL3, LHX1, FOXC1) (Fig. 3D, Fig. 3E) or non-mesodermal fate (fig. S3A). At d4.5 and d7.5, KO and HAase-treated cardioids continued to show the downregulation of cardiac/muscle-related pathways, while suggesting misspecification into non-cardiac cell fates (Fig. 3F, fig. S3D, fig. S3E, fig. S3F, fig. S3G, fig. S3H, fig. S3I). At these later stages, KO and HAase-treated cardioids downregulated genes that respond to muscle stretch (e.g. ANKRD1, NPPA, NPPB, MYH7, CSRP3) (Fig. 3G). At d7.5, both KO and HAase-treated samples showed aberrant regulation of genes involved in cell cycle and metabolic processes (fig. S3G, fig. S3H, fig. S3I). Overall our genetic and enzymatic perturbations in cardioids demonstrate that HA is required for a timely transition of nascent mesoderm into pre-cardiac mesoderm and cardiomyocytes.

**Figure 3.**
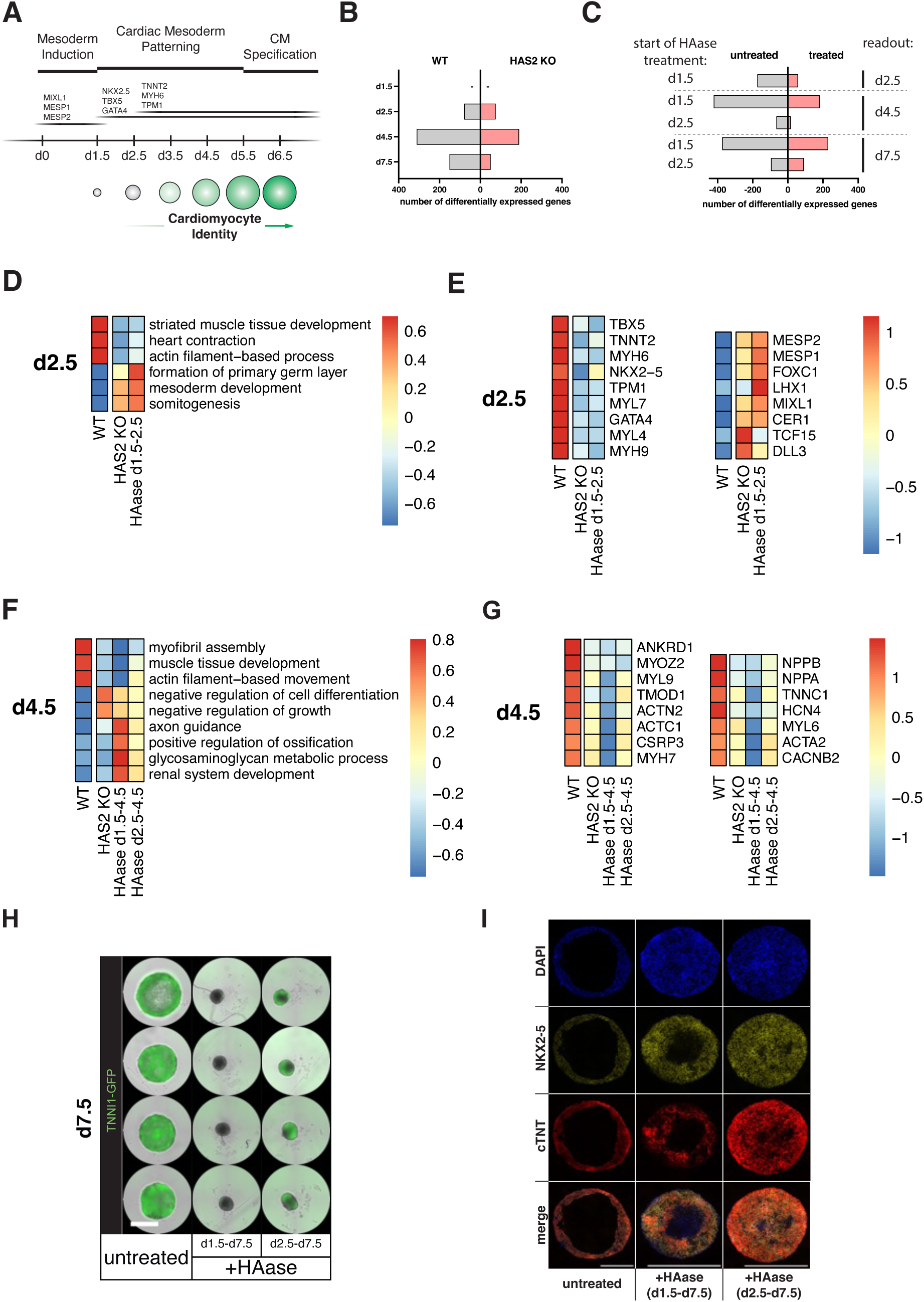
HAS2 and hyaluronan are necessary for cardiac mesoderm specification. (A) Timeline of cardioid fate specification and expression of exemplary regulatory and structural genes. (B) Number of differentially expressed genes of WT vs. HAS2 KO cardioids at different readout timepoints. (C) Number of differentially expressed genes of untreated vs. HAase-treated cardioids (treatment either started at d1.5, or d2.5) at different readout timepoints. (D) Enrichment scores of selected pathways differentially regulated between WT, HAS2 KO and HAase-treated cardioids at d2.5. See fig. S3A for complete list. (E) Expression of selected genes differentially regulated between WT, HAS2 KO and HAase-treated cardioids at d2.5. (F) Enrichment scores of selected pathways differentially regulated between WT, HAS2 KO and HAase-treated cardioids at d4.5. See fig. S3D for complete list. (G) Expression of selected genes differentially regulated between WT, HAS2 KO and HAase-treated cardioids at d4.5. (H) Images showing brightfield and TNNI-GFP signal of untreated vs. HAase-treated cardioids at d7.5. Scale bar 1000 microns. (I) Immunostaining of cryosections of untreated vs. HAase-treated cardioids at d7.5 showing absence of cavities in treated cardioids. Note lower differentiation efficiency in samples treated from d1.5 (as compared to samples treated from d2.5). Scale bar 500 microns. Color bars in (D) and (F): GSVA enrichment score. Colors bars in (E) and (G): Scaled gene expression.

To define the precise temporal requirement of HA in cardioid development, we compared HAase treatment initiated at d1.5 versus d2.5 (Fig. 3C). Direct comparison of these two conditions at d4.5 shows that the aforementioned effect is weakened in cardioids treated only from d2.5 on (Fig. 3F, Fig. 3G, fig. S3D, fig. S3E, fig. S3F, fig. S3G, fig. S3H, fig. S3I), suggesting that the presence of HA from d1.5 is critical for instructing cardioid development. To complement our transcriptomic experiments, we studied the expression of cardiac proteins in cardioids treated with HAase. HAase treatment from d1.5 largely reduced TNNI1-GFP expression at d7.5, while delaying the treatment to d2.5 reversed this effect in similarly size-reduced cardioids (Fig. 3H). While cells co-express NKX2-5 and cTNT by d7.5 in untreated cardioids, we found many NKX2-5-positive cells that failed to co-express cTNT in cardioids treated with HAase from d1.5 (Fig. 3I). Taken together, we identify d1.5 to d2.5 as the critical time window in cardioid development during which the presence of HA hydrogel is required for the proper specification of the cardiac mesoderm and subsequent formation of cardiomyocytes.

### Loss of hyaluronan compromises the beating function of left ventricular cardioids

After finding that HA underlies morphogenesis and cell fate specification, we asked whether HA plays a role in cardioid physiology. The earliest contractions in LV cardioids begin around d5, with all samples showing consistent beatings by d5.5(*14*) (Fig. 4A, 4B). At d5.5, cardioids generated from HAS2 KO cell lines showed an overall reduced beating frequency compared to WT cardioids (KO3 showed no contractions) (Fig. 4B’). The difference in beating frequency for WT vs KO cell lines became statistically significant at day 6.5 and 7.5 (fig. S4A). Starting from day 5.5, contractile HAS2 KO cardioids (KO1, KO2) showed a significantly reduced contraction amplitude, which can be described as a ‘twitching’ behavior as opposed to the large amplitude beating of WT cardioids (Fig. 4B, Fig. 4B’’, fig. S4B, Movie S6). Overall, reductions in beating frequency and contraction amplitude functionally distinguishes HAS2 KO from WT cardioids.

**Figure 4.**
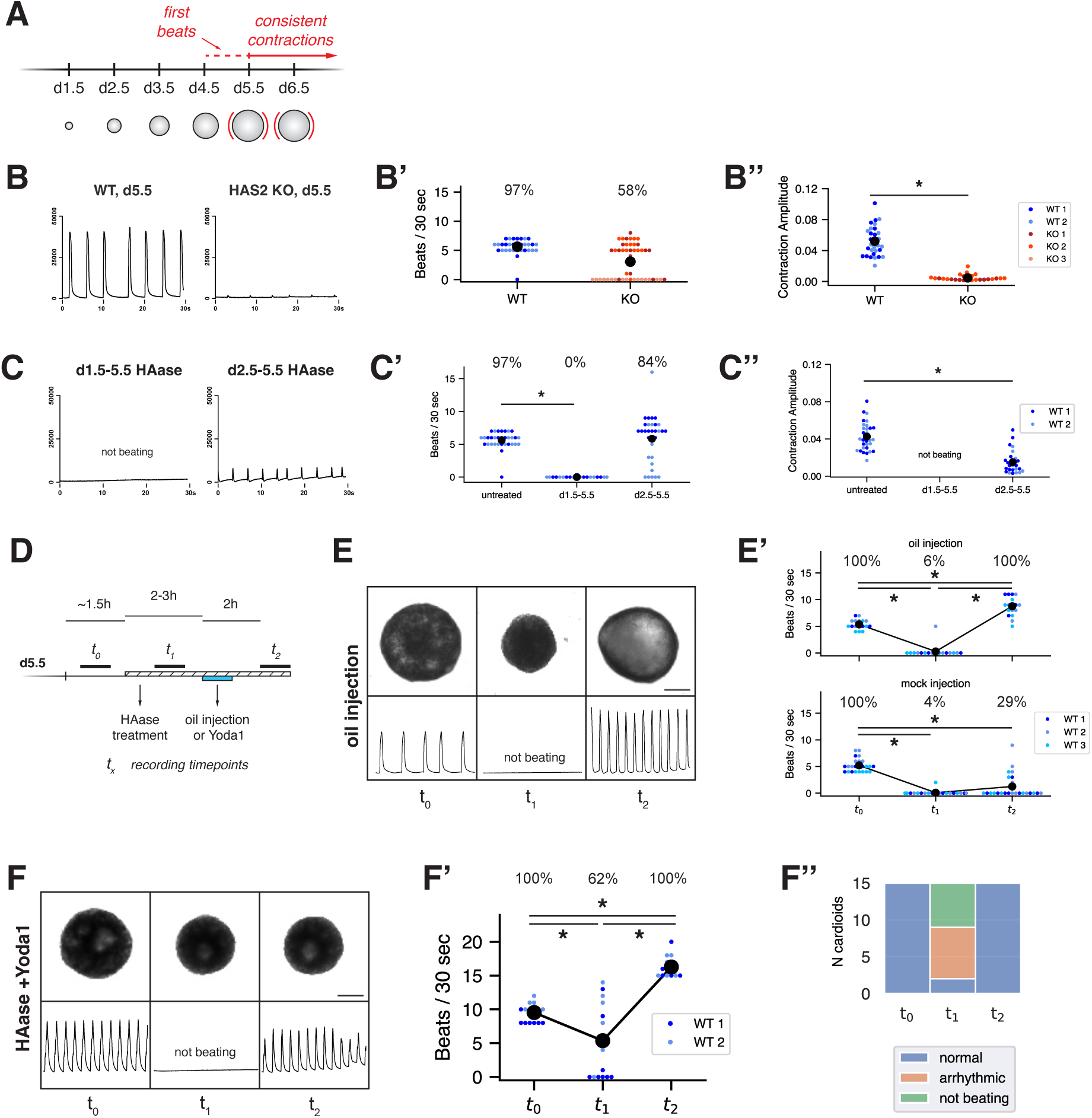
Hyaluronan provides a physical environment that is necessary and sufficient for the beating function of LV cardioids. (A) Schematic of LV cardioid development highlighting the onset of contractile activity. (B) Contraction traces of day 5.5 WT and HAS2 KO cardioids. (B’) Beating frequency and (B’’) Contraction Amplitude of day 5.5 WT and HAS2 KO cardioids. Colors distinguish samples prepared from n=2 WT cell lines and n=3 HAS KO cell lines. n = 16 samples per cell line per condition. (C) Contraction traces of day 5.5 cardioids treated with HAase for different time windows. (C’) Beating frequency and (C’) Contraction Amplitude of day 5.5 cardioids treated with HAase for different time windows. Colors distinguish samples prepared from two cell lines. n=16 samples per cell line per condition. (D) Schematic of experiment to test the effect of acute hyaluronic acid degradation and mechanical rescue strategies on the beating function of day 5.5 cardioids. (E) Bright field image (top) and contraction traces (bottom) of a day 5.5 cardioid subjected to HAase treatment, followed by oil injection. Scale bar 500 microns. (E’) Top: Beating frequency of cardioids subjected to HAase treatment, followed by oil injection. Bottom: Beating frequency of cardioids subjected to HAase treatment, followed by mock injection. Colors distinguish samples prepared from three cell lines. n=5-8 samples per cell line per condition. (F) Bright field image (top) and contraction traces (bottom) of a day 5.5 cardioid subjected to HAase treatment, followed by Yoda1 treatment. Scale bar 500 microns. (F’) Beating frequency of cardioids treated with HAase and Yoda1. (F’’) Classification of arrhythmia phenotypes. n=15 samples from two cell lines are combined. See also fig. S5D. Percentages indicate the fraction of beating samples in each condition. Asterisks indicate a statistically significant difference of p-value < 0.05.

To define the temporal HA requirement for contractile function, we treated cardioids with HAase for different time windows and assessed contraction at d5.5 and 6.5 (fig. S4C). When the treatment started at d2.5, beating frequency of the treated group was comparable to the control group at d5.5 (Fig. 4C’), but most cardioids stopped beating by d6.5 (fig. S4D). Contraction amplitude was significantly reduced relative to untreated samples (Fig. 4C’’, fig. S4E, Movie S7), similar to the ‘twitching’ phenotype observed in HAS2 KO cardioids. When HAase treatment started at d1.5, this completely abolished contractions in cardioids (Fig. 4C, Fig. 4C’, fig. S4D, Movie S7). In summary, our functional experiments revealed that HA is particularly important during the d1.5-d2.5 developmental window, when it contributes to the beating frequency and contraction amplitude to ensure proper LV cardioid function at later stages.

### Hyaluronan provides a physical environment that is necessary and sufficient for the beating function of left ventricular cardioids

One plausible explanation of impaired beating of LV cardioids devoid of HA from d1.5 is that the cells fail to differentiate into functional cardiomyocytes (see Fig. 3). An additional possibility is that from d5.5, the physical properties of HA hydrogel contribute to the normal beating of LV cardioids. To directly test this scenario, we treated beating LV cardioids at d5.5 with HAase and immediately assessed beating (Fig. 4D). At 1 hour post HAase treatment, cardioids were reduced in size (Fig. 4E, fig. S4F) and showed a near complete loss of contractility (Fig. 4E, Fig. 4E’ at *t_1,_* Movie S8), indicating that acute HA degradation impairs cardioid beating. To ask if this beating defect can be physically rescued in the absence of HA, we manipulated tissue morphology and mechanics by injecting an inert oil droplet into the collapsed cavities of HAase-treated cardioids (Fig. 4D). Oil injections successfully increased cardioid size with diameters approaching values of untreated samples (Fig. 4E, fig. S4F at *t_2_*). Strikingly, oil-injected cardioids restored their beating, while mock-injected remained largely inactive (Fig. 4E, Fig. 4E’ at *t_2,_* Movie S8). Taken together, our results demonstrate that HA provides a physical environment that is essential for the normal beating of LV cardioids. However, these experiments did not address whether this physical environment is defined by tissue morphology or mechanics.

### Mechanosensitive channels link hyaluronan-generated forces to left ventricular cardioid beating

Cardiomyocytes are mechanosensitive cells that detect and respond to physical forces through multiple molecular mechanisms (*28*). Key among these are mechanosensitive channels (MSCs) that translate tissue-scale forces to cellular electrophysiological responses (*29*). To test whether MSCs underlie cardioid contractility, we used the MSC inhibitor GsMTx-4, which broadly targets the Piezo and TRP families of mechanosensitive channels (*30*, *31*). Treating d5.5 LV cardioids with 5.0 µM GsMTx-4 abolished contractions (fig. S4G, fig. S4H, Movie S9), but had no effect on tissue size (fig. S4G, fig. S4I). At a lower dose of 2.5 µM GsMTx-4, we observed that cardioids from one WT cell line continued to beat (fig. S4H, WT3), yet, these contractions were predominantly arrhythmic (fig. S4J, fig. S4K, fig. S4L, Movie S9). To ask if activation of a specific MSC results in cardioid contractions, we turned to the Piezo1 activator Yoda1 (*32*, *33*) (Fig. 4D). Treating LV cardioids with Yoda1 increased the beating frequency (fig. S4M, Movie S10). Strikingly, Yoda1 treatment completely rescued the HAase-induced beating defects, including arrhythmia (Fig. 4F, Fig. 4F’, Fig. 4F’’, fig. S4N, fig. S4O, Movie S10), despite the Yoda1-treated cardioids remaining small in size (Fig. 4F, fig. S4P). Taken together our pharmacological manipulations suggest that it is not the HA-induced morphology *per se* but the mechanical forces generated by HA that act through MSCs and prime LV cardioids for rhythmical contractions.

### Multi-chamber architecture provides resilience to mechanical perturbation *in vivo* and in vitro

Finally, we asked whether the mechanical role of HA in early cardiac contractions is a general phenomenon beyond LV cardioids. To test this, we chose chick embryos as an accessible source for whole functional hearts. HAase injection into cultured chick embryos at HH8 increased the volume of the heart lumen at the expense of cardiac jelly as reported previously (*34*) (fig. S5A). In some cases, the volume of the heart tube expanded (*26*, *35*) (fig. S5B), potentially caused by systemic blood pressure. Therefore, to study the beating function of whole hearts in mechanical isolation from the embryo, we explanted hearts from HH11-12 chick embryos and cultured them *ex vivo*. HAase treatment readily reduced the size of heart explants, while the size of control samples remained unchanged (Fig. 5A, Fig. 5A’, fig. S5C). However, the beating frequency was not significantly affected with or without HAase treatment (Fig. 5A’’, fig. S5D, Movie S11). This indicates that the chick embryonic hearts, like those of zebrafish(*36*), are functionally resilient to the acute degradation of HA.

**Figure 5.**
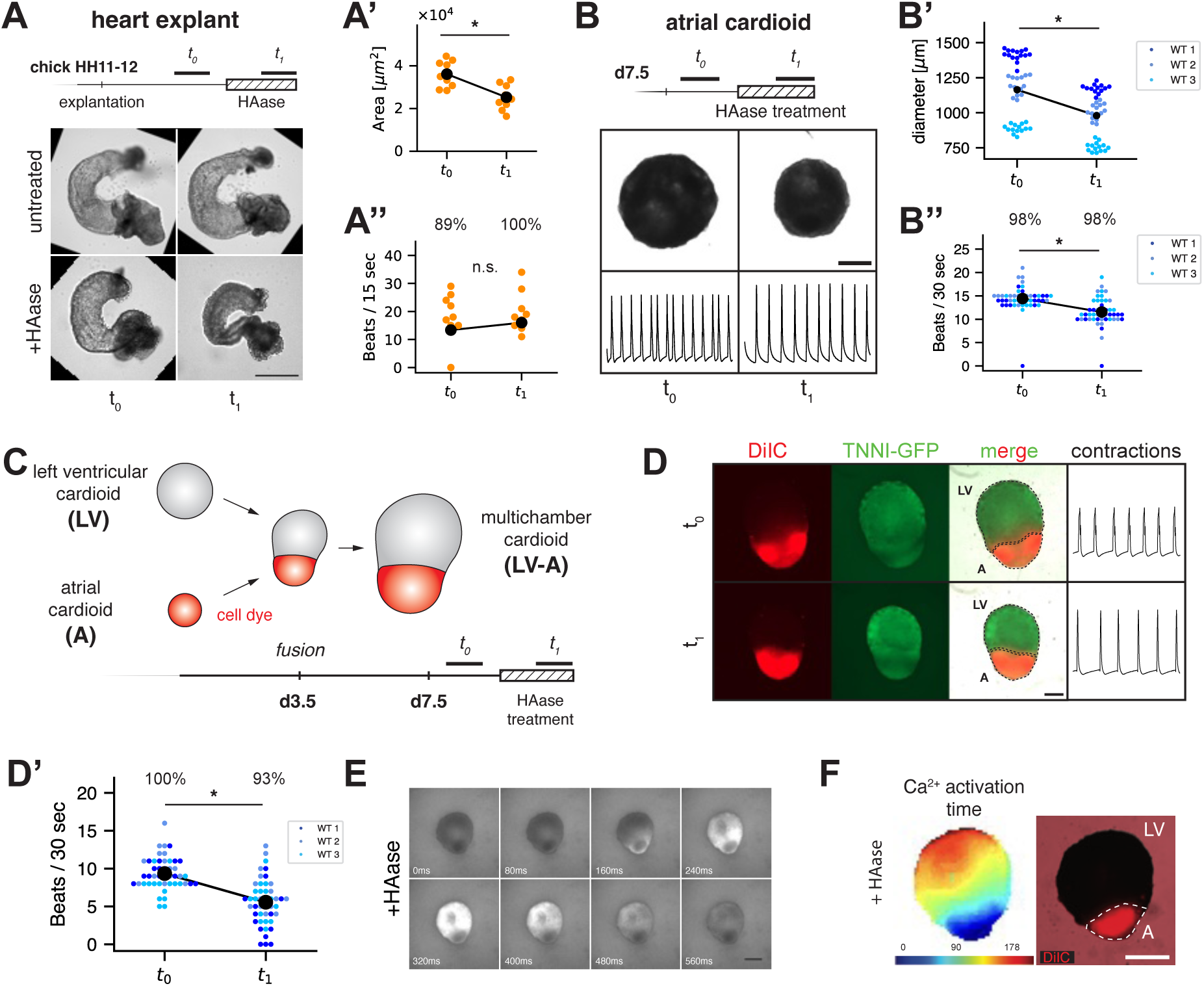
Multi-chamber architecture provides resilience to mechanical perturbation *in vivo* and *in vitro*. (A) Experiment to test the effect of acute HA degradation on chick heart explants. Each row of the image panel shows the same heart explant before and after hyaluronidase treatment. t1 recording was done 4.3 hours after addition of HAase. Scale bar 500 microns. (A’) Area and (A’’) Beating frequency of heart explants before and after HAase. n=9 samples. (B) Experiment to test the effect of acute HA degradation on day 7.5 atrial cardioids. Brightfield images and contraction traces of the same cardioid before and after HAase treatment. t1 recording was done 1 hour after addition of HAase. Scale bar 500 microns. (B’) Diameter and (B’’) Beating frequency of atrial cardioids before and after HAase treatment. Colors distinguish samples prepared from three cell lines. n=13-16 samples per cell line per condition. (C) Schematic of experiment to assay the role of acute hyaluronic acid degradation on multi-chamber cardioids. LV-A multi-chamber cardioids are generated by fusing an atrial cardioid (labelled with a dye) and a left ventricular cardioid at day 3.5, allowed to develop, and assayed at day 7.5. t1 recording was done 1 hour after addition of HAase. (D) Brightfield and fluorescent images of the same day 7.5 LV-A multi-chamber cardioid before and after HAase treatment. TNNI-GFP is expressed in both chambers, while only the atrial chamber is labelled with the dye DilC18(5)-DS. Contraction traces are shown for the corresponding timepoints. Scale bar 500 microns. (D’) Beating frequency of LV-A multichambered cardioids before and after HAase treatment. Colors distinguish samples prepared from three cell lines. n=14-16 samples per cell line per condition. (E) Ca^2+^ signal propagation in GCaMP6f-expressing LV-A cardioids after HAase treatment. Scale bar 500 microns. (F) Left: Heatmap indicates the time when Ca^2+^ signal first appeared. Right: DilC18(5)-DS signal indicates the region corresponding to the atrial chamber. Scale bar 500 microns. Percentages indicate the fraction of beating samples in each condition. Asterisks indicate a statistically significant difference of p-value < 0.05.

Unlike LV cardioids, the chick heart contains multiple heart regions, including the primitive atrium, which acts as the pacemaker region prior to the development of the sinoatrial node (*37*, *38*). To dissect the region-specific functions of the human heart, we generated atrial cardioids (A) and treated them with HAase. We performed our experiments at d7.5, a timepoint when all A cardioids show robust beating. HAase treatment reduced cardioid size, suggesting that HA supports cavity morphogenesis of A cardioids (Fig. 5B, Fig. 5B’) similar to LV cardioids. However, the beating frequency of A cardioids decreased only slightly (Fig. 5B’’). Thus, in contrast to LV cardioids, A cardioids are functionally resilient to mechanical unloading. Our findings in chick explants and A cardioids led us to ask if a multichamber architecture is sufficient to confer functional resilience to mechanical unloading. To this end we fused LV and A cardioids (*15*) at d3.5 and treated them with HAase at d7.5 (Fig. 5C). Untreated LV-A multichamber cardioids showed a beating frequency of 8.7±2.6 beats/30s, which is between the frequencies of isolated A (14.7±1.6 beats/30s) and LV (1.6±0.9 beats/30s) cardioids at this day (Fig. 5D, Fig. 5D’). HAase treatment reduced the area of LV-A multichamber cardioids by 27% (fig. S5E) and led to a decrease in beating frequency (Fig. 5D’ at *t_1_*). However, unlike LV cardioids, the majority of LV-A multichamber cardioids continued beating after HAase treatment. Brightfield movies confirmed contraction in both A and LV regions (Movie S12). To better understand the spatial and mechanistic basis of contractions in these conditions, we performed optical mapping of Ca^2+^-transients using cardioids expressing GCaMP6f (*39*). Upon HAase treatment, LV cardioids lost their Ca^2+^-transients, while they were maintained in A cardioids, consistent with their contractilities in the respective conditions (Movie S13). In multichamber cardioids, Ca^2+^-transients initiated in the atria and propagated to the ventricular portion in all cases for untreated and HAase-treated conditions, respectively (Fig. 5E, fig. S5F), indicating that atrial pacing can rescue the beating defect of LV cardioids. Altogether, we show that cardioid sub-compartments exhibit distinct mechanosensitivity and that the multichamber architecture extends functional resilience across distinct heart chambers.

## DISCUSSION

Organ shape and function emerge from the dynamic interactions between differentiating stem cells and microenvironmental cues. In this study, we found that human cardioids endogenously produce a hyaluronic acid (HA) hydrogel that plays a fundamental role in the development of cardiac form, fate, and function. Within two days of cardioid development, progenitor cells begin to express the hyaluronic acid synthase, HAS2, which is responsible for producing the HA hydrogel that swells and drives cavity formation (Fig. 1, Fig. 2). HA-enriched cavities occupy up to 40% of the total volume of cardioids, contributing to rapid volumetric growth. In contrast to the widespread mode of lumen formation that involves active ion transport in polarized cellular sheets (*18–21*), cavity formation by hydrogel swelling creates an acellular space, accessible to large, freely diffusing molecules in a porous tissue. The HA hydrogel in cardioids is reminiscent of the cardiac jelly in the early vertebrate heart. The HA secreted by the cardiac mesoderm and cardiomyocyte progenitors via HAS2 is the major component of the cardiac jelly, driving the expansion of the myoendocardial space (*23–26*, *34*). Thus, using cardioids, we uncover and dissect a general biophysical mechanism that rapidly generates acellular space in developing tissues (*40*, *41*), providing the first mechanistic evidence for the role of HA in human cardiac morphogenesis.

Beyond its role in sculpting cardioid chambers, we found that HA is crucial for hierarchical specification of cardiac fate from nascent mesoderm (Fig. 3). Genetic and enzymatic degradation of HA led to aberrant expression of presomitic mesoderm genes and attenuated cardiac lineage commitment in cardioids. We established that HA mainly acts during a narrow developmental window (d1.5-2.5), and propose three mechanistic scenarios for the role of HA in cell fate regulation. First, an aberrant metabolism in HAS2 knockout cells (*42*, *43*) and conditions with high hyaluronidase activity (*44*), resulting from disrupted UDP-sugar utilization or altered glycolytic flux, may instruct the cell cycle and progenitor lineage decisions (*45*, *46*). Second, HA receptor-mediated signaling through receptors (e.g. CD44 (*47*)) may activate downstream transcriptional networks for cardiac differentiation (*25*). Third, mechanical forces generated by the HA hydrogel may directly regulate gene expression programs required for cardiac specification (*48*). Future studies using cardioids are well-positioned to dissect these scenarios to understand cell fate decisions at the onset of human cardiogenesis.

While the effects of ECM components on cardiac differentiation and cell fate had been studied (*10*, *11*), cardioids uniquely enabled the investigation of early functional consequences, revealing a purely mechanical role of HA for cardiac contraction in a human model (Fig. 4). Based on our acute perturbation and rescue experiments, we propose that the swelling forces generated by HA hydrogel establish a permissive mechanical condition. We speculate that HA stretches the cardiomyocytes, mimicking the state of the heart under preload, supporting robust contractions in left ventricular cardioids. Our findings extend the well-established mechanosensitive nature of cardiomyocytes (*49*, *50*) by demonstrating that these cells actively synthesize ECM components to engineer their own mechanical microenvironment for optimal contraction. We identified the mechanosensitive ion channel Piezo1 as a crucial mediator of ECM-derived swelling force and cellular response. Future studies will leverage the experimental accessibility of cardioids to address how ECM-derived mechanical signals are transmitted through changes in membrane potential and Ca^2+^ signaling to ultimately regulate the earliest contractions of the heart (*51*).

We generated cardioids corresponding to different heart regions and discovered chamber-specific mechanosensitivity, achieving a level of functional dissection previously unattainable (Fig. 5). Strikingly, in contrast to left ventricular cardioids, the contraction of atrial cardioids showed resilience to HA removal. When integrated into multi-chambered cardioids, the atrial portion drives the contractions in the left ventricular cardioid even after hyaluronidase treatment. The resilience of multi-chamber cardioids mirrors the mechanical robustness of whole heart explants. In the human embryo, the first heartbeat occurs around 20 days post-fertilization and is thought to originate from the region corresponding to the future left ventricle. As the atrial region develops, it assumes the role of pacemaker with increasingly consistent tempo. Left ventricular cardioids develop contractility two days earlier than atrial cardioids (*15*), but whether regional differences in ECM production (*52*) contribute to this accelerated developmental timing and pacemaking remains unexplored. We speculate that the complementary chamber-specific characteristics in ECM profile, mechanosensitivity, and developmental timing ensure a robust and timely emergence of organ-scale physiology in multi-chambered hearts.

Unlike exogenous matrices that dominate current organoid culture and tissue engineering approaches (*53*, *54*), endogenous cell-synthesized ECM provides a dynamic, physiologically relevant environment for coordinating form, cell fate, and tissue function. Our findings indicate that cardiac tissue generation from human PSCs should incorporate mechanical cues as early as day 1.5 of differentiation. This contrasts with most current methods for generating human cardiomyocytes (*55*), which employ 2D differentiation conditions for multiple weeks before replating, scaffolding, or bioprinting for functional analysis. Our left ventricular cardioid protocol achieves high cardiomyocyte differentiation efficiency (*14*) and robust beating by day 5.5, representing one of the fastest timelines from hPSCs to contractile tissue. We attribute this efficiency to early 3D self-organization of cardiac progenitor cells that capture endogenous cell-secreted HA for efficient cardiac lineage commitment. Furthermore, the accumulated HA establishes a favorable physical environment for robust contraction. Consistent with this principle, other studies differentiating human PSCs at high density similarly report chamber formation and accelerated contractility (*16*, *17*, *56*). These findings collectively suggest that recapitulating the mechanical microenvironment during early organogenesis may be a generalizable strategy to accelerate the generation of more functional and physiologically relevant tissues. More broadly, by understanding and rewiring cellular programs for dynamic ECM synthesis, we can better control form, fate, and function to engineer advanced organoids.

### Limitations and Future Directions

While cardioids offer a powerful modular platform for studying cardiac biology, their simplicity and our current understanding impose important limitations. As cardioids do not incorporate mechanical input from the foregut endoderm, this model may mimic heart formation in *cardia bifida* embryos, where foregut constriction is abolished (Fig. S5G, Fig. S5H). Moreover, we cannot currently direct HA generation between myocardial and endocardial tissue layers to reproduce the tube-within-tube architecture of the developing heart. Thus, we still lack directional flow or associated mechanical cues like wall shear forces as well as direct measurements of hydrogel rheology and tissue-scale forces. New technologies for *in situ* mechanical actuation and measurements may overcome this gap. Finally, the physicochemical profile of the ECM in cardioids remains unknown. Characterizing the molecular weight of HA and the presence of other ECM components is essential for understanding how this ECM is dynamically remodeled and sensed by cells. Such insights may enable dissection of later developmental events like trabeculation and valvulogenesis in this tractable model of the human heart.

## ACKNOWLEDGEMENTS

We thank all members of the Mendjan and Ishihara labs for their help and discussions. We thank Katarzyna Warczok and Megan Shi for lab management. We are grateful for IMBA/IMP/Vienna Bio Center core facilities (Bioinformatics (Maria Novatchkova), BioOptics, Molecular Biology Service, Electron Microscopy, Histology, Next Generation Sequencing, and Stem Cell Core facilities) for their services. We thank the Allen Institute and HeartBeat.bio for cell lines.

## Funding

This work was supported by the Austrian Academy of Sciences (OEAW), an ERC Advanced Grant (CardioGROWTH, 101201790) to S.M.; an Austrian Science Fund (FWF) grant (ID: PAT4435223) to S.M.; an American Heart Association award (25CDA1432232) to K.I.; and a NIH grant (R01GM134226) to A.R.H.

## Author contribution

S.M.J., K.I., and S.M. conceived the project. S.M.J. performed all experimental work with assistance from others. A.D. contributed to multi-chamber cardioid experiments, mechanosensitive perturbations, and calcium imaging. J.K. generated and assayed HAS2 knockout cell lines and hyaluronidase experiments. V.G. performed computational analysis of RNAseq data. Y.K. performed computational analysis of 3D cavity morphology. M.M. assisted with experiments and image analysis. T.I. generated CAG-GFP and FUCCI cell lines. K.I. performed 3D imaging of cleared samples and lightsheet imaging.

K.I. performed all quantitative analysis and statistics except RNAseq analysis. S.M.J., K.I., and S.M. wrote the manuscript with input from all authors. K.I. and S.M. supervised the work and obtained funding.

## Competing interests

S.M. is a co-founder and scientific advisory board member of Heartbeat Bio, a IMBA spin-off; IMBA filed patent applications with S.M (EP4540370A1) and S.M.J. and S.M. as co-inventors (EP4483888A3).

## Data, code and materials availability

RNA-seq data has been deposited to the NCBI Gene Expression Omnibus and is accessible through the GEO accession number GSE317879. 3D imaging data will be deposited in the BioImage Archive. Custom analysis code is available from the corresponding authors upon reasonable request. Cell lines and plasmids are available upon reasonable request from the corresponding authors and may require a material transfer agreement.

## LIST OF SUPPLEMENTARY MATERIALS

### Supplementary Materials

Materials and Methods

Figs. S1 to S5

Tables S1 and S2

Movies S1-S13

## SUPPLEMENTARY FIGURE LEGENDS

**Fig. S1.**
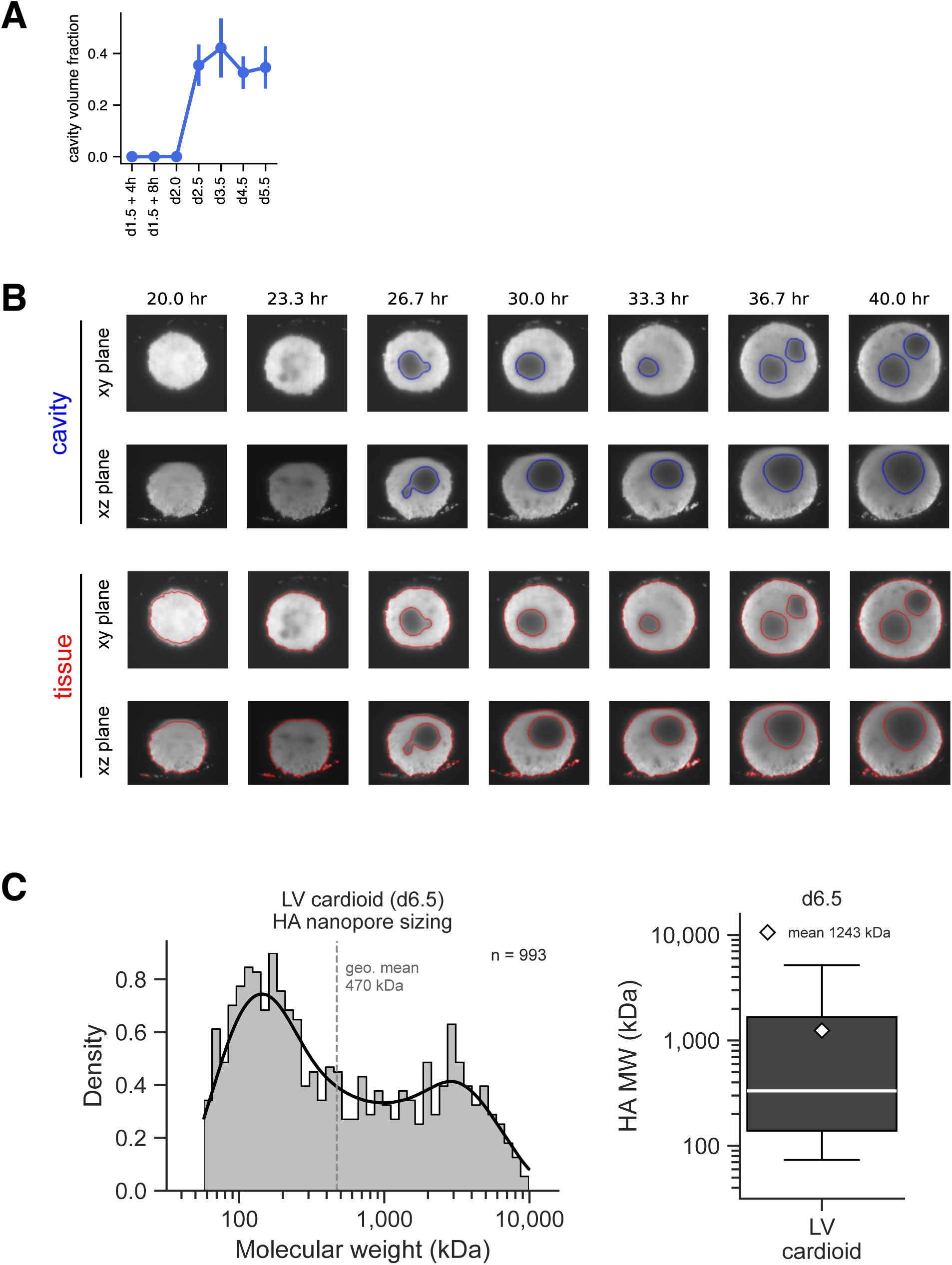
(A) Cavity volume fraction, defined as the total cavity volume over total cardioid volume, of the samples in Fig. 1C. Error bars indicate the standard deviation across n=4-12 samples per time point. (B) Cavity and cell-containing tissue regions are segmented from the light sheet microscopy data for the sample in Fig. 1D. (C) Molecular weight distribution of HA extracted from day 6.5 LV cardioids, quantified by solid-state nanopore analysis.

**Fig. S2.**
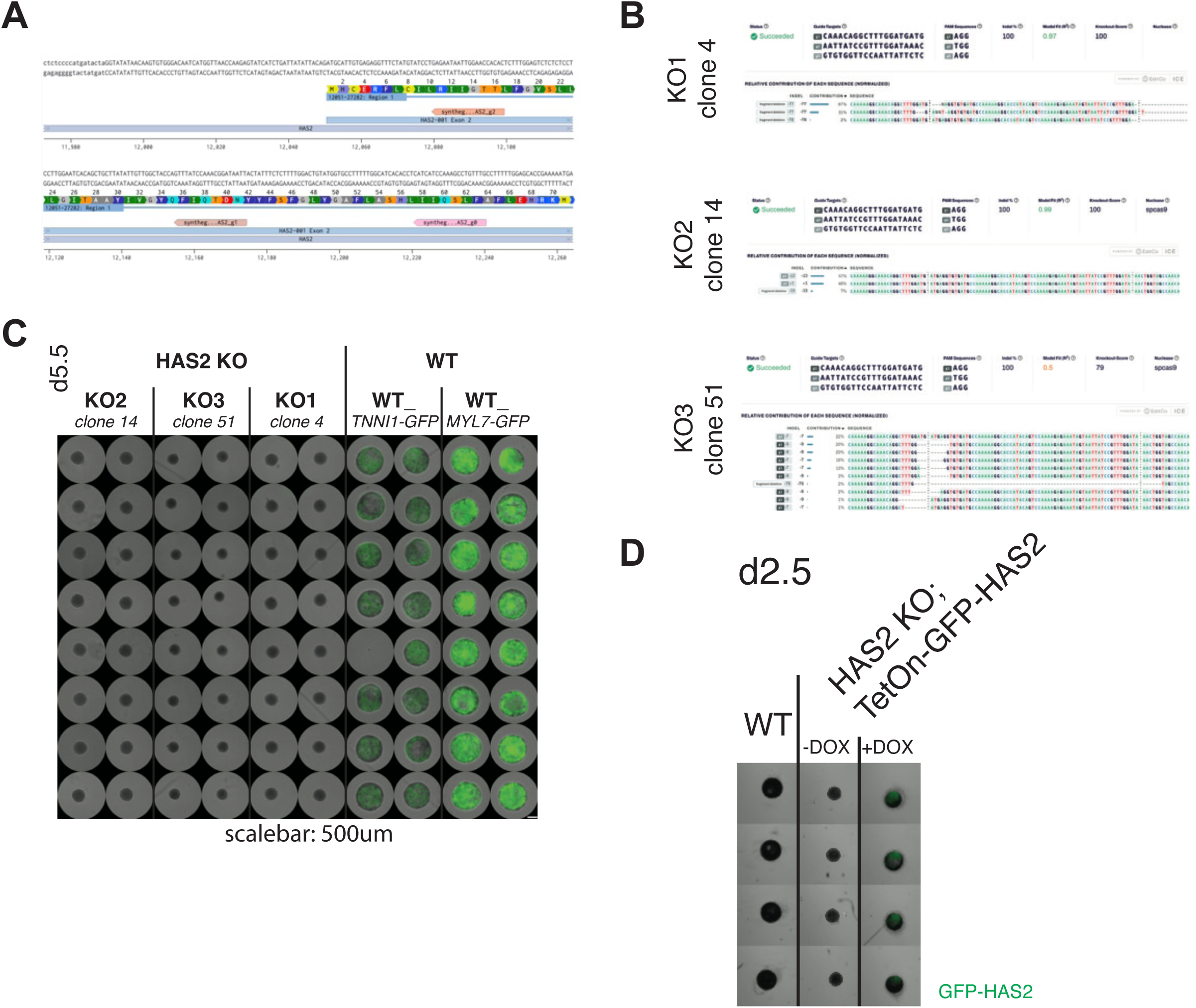
(A) Schematic of gRNA design to knock out HAS2 gene in human PSCs. All three gRNAs (g0, g1, and g2) were used simultaneously. (B) Insertion-Deletion analysis of selected HAS2 KO clones used in this study. (C) Brightfield images of d5.5 cardioids generated from HAS2 KO cell lines (KO1, KO2, and KO3) and HAS2 WT lines (TNNI-GFP and MYL7-GFP). (D) Brightfield and fluorescent images of WT and HAS2 KO; TetOn-GFP-HAS2 cardioids at day 2.5. Doxycycline was added to the indicated condition at day 1.5. Same experiment as Fig. 2F.

**Fig. S3.**
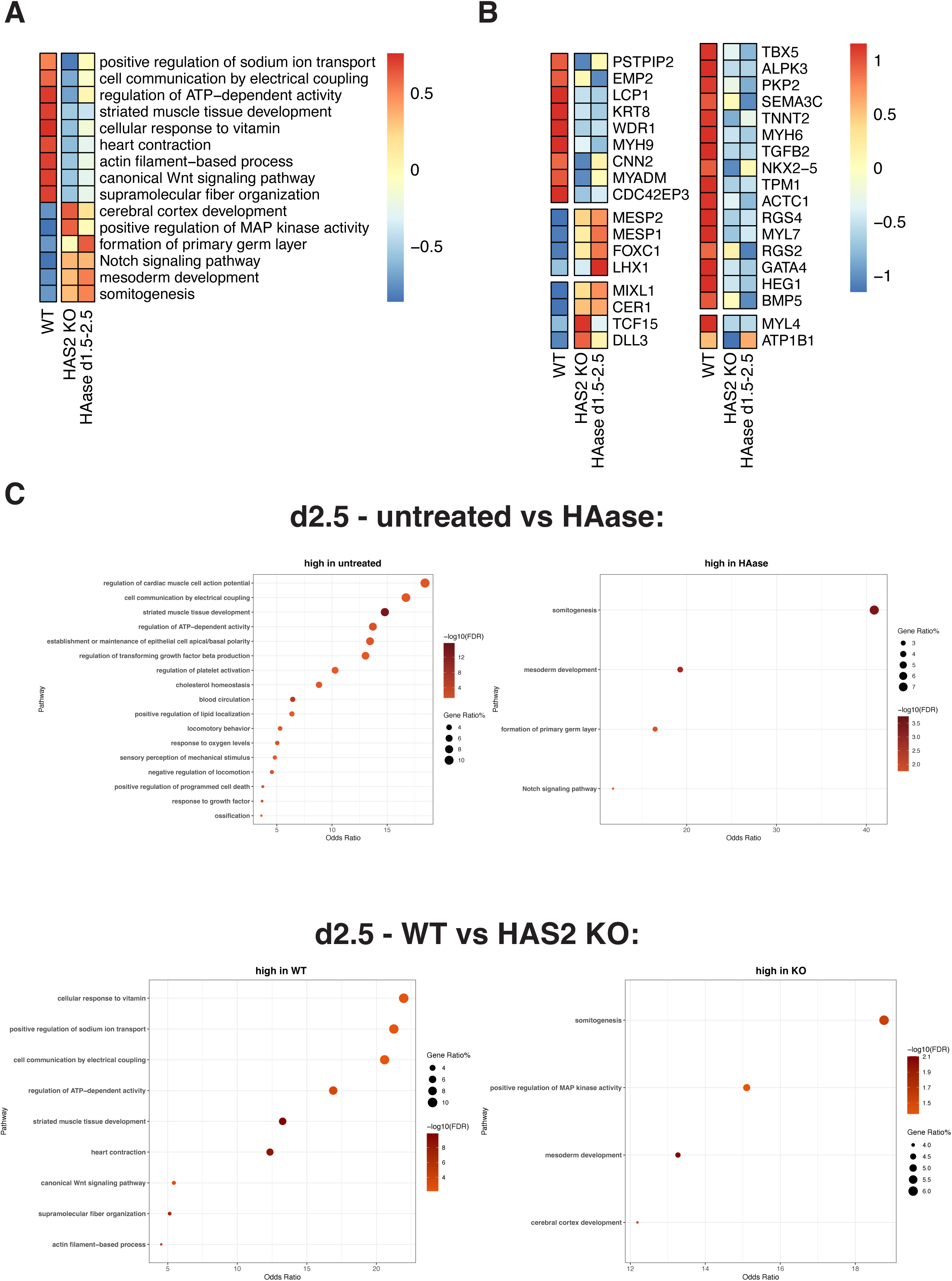
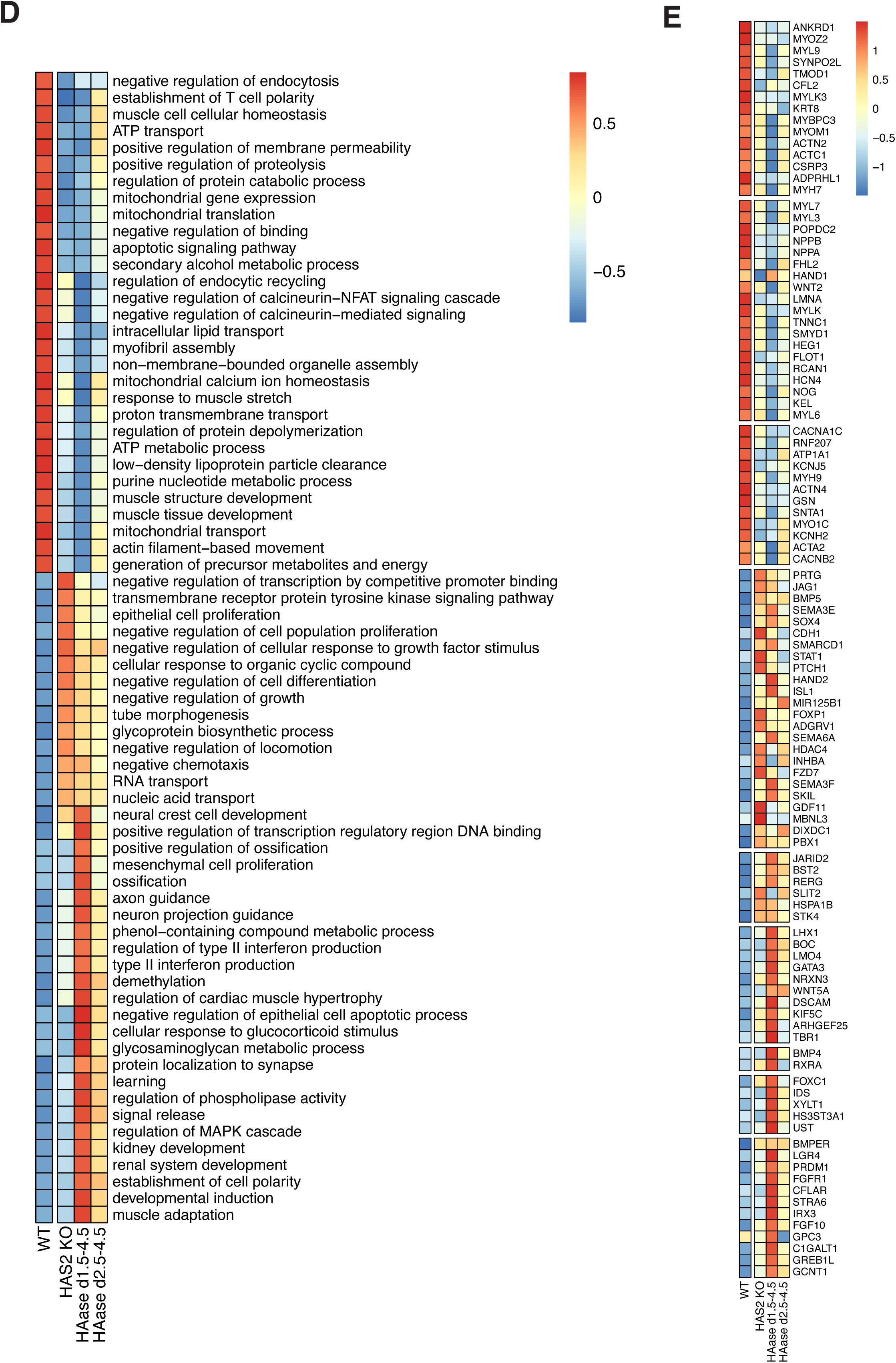
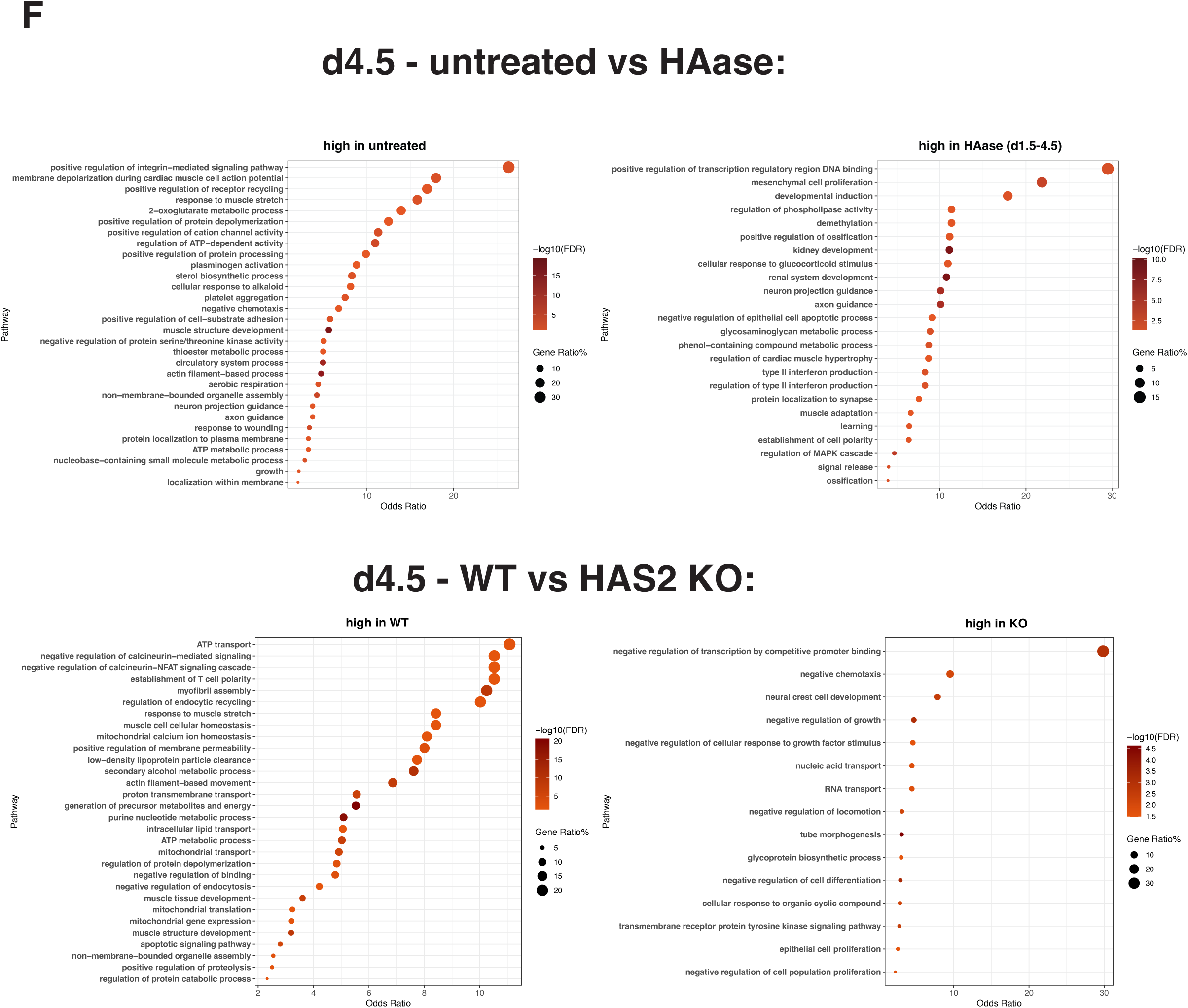
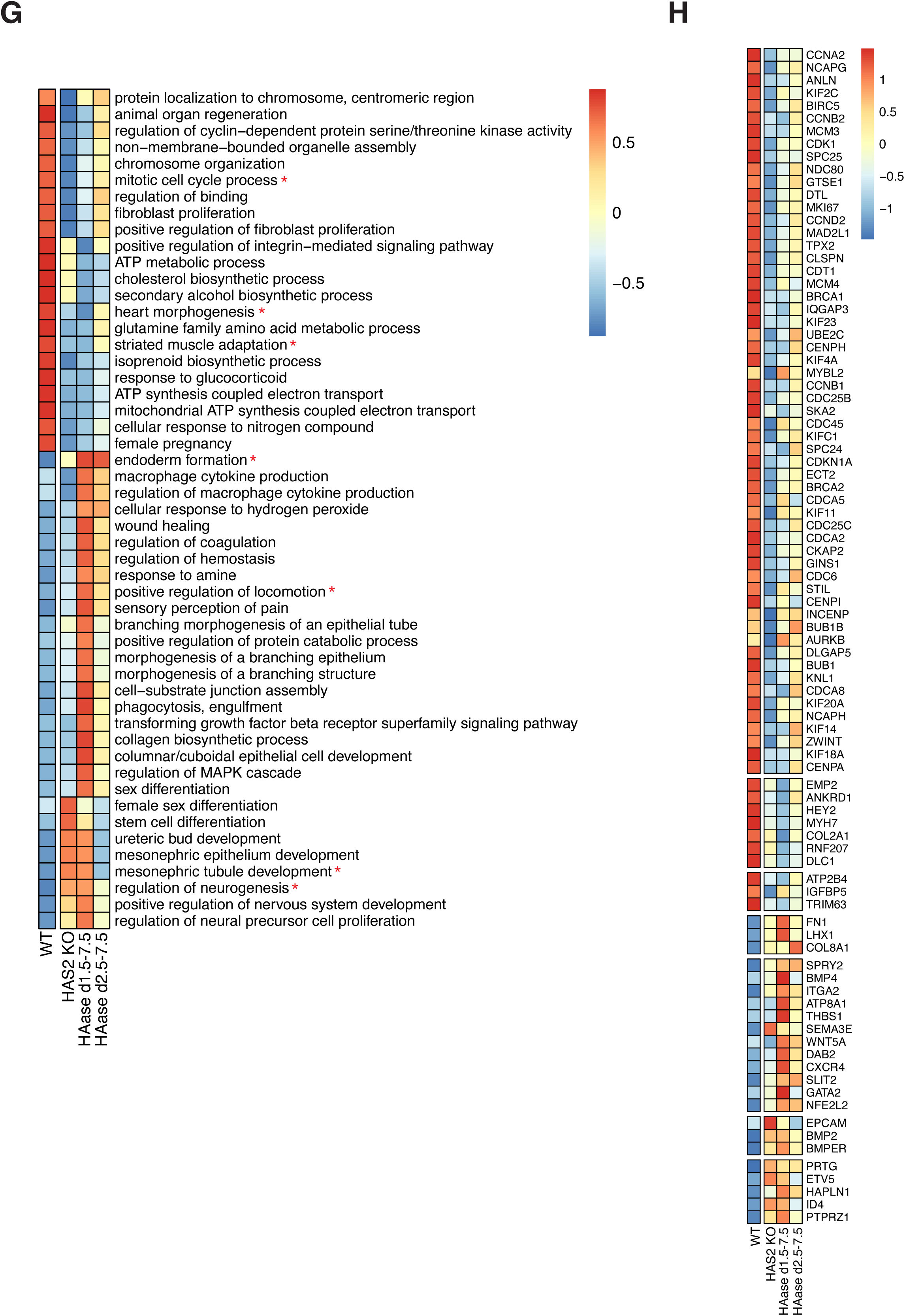
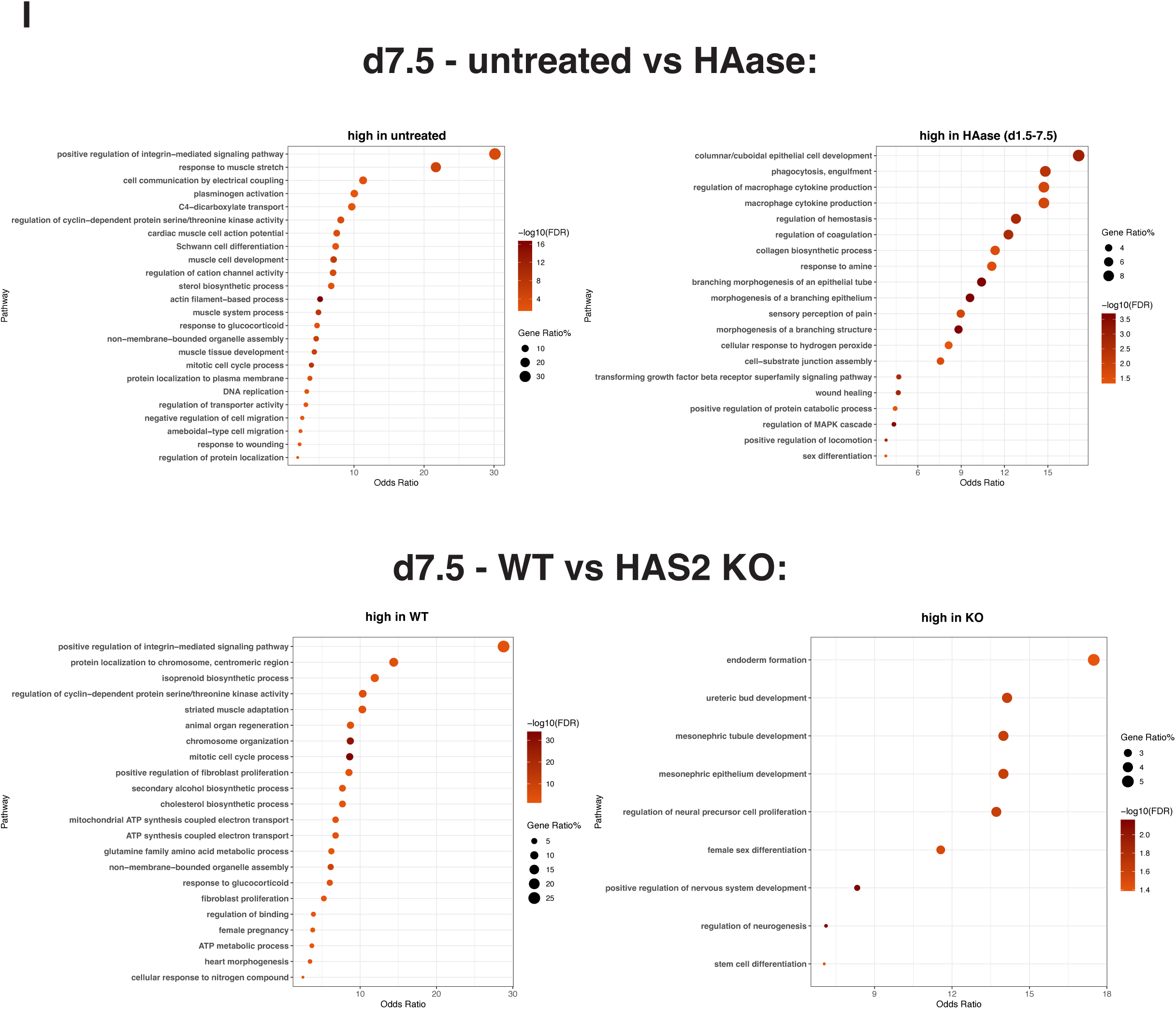
(A) Complete list of pathways differentially regulated between WT, HAS2 KO and HAase-treated cardioids at d2.5. (B) Expression of genes representing pathways shown in Fig. 3D. (C) Dot plots indicating GO Biological Process Term enrichment for the indicated comparisons at d2.5. (D) Complete list of pathways differentially regulated between WT, HAS2 KO and HAase-treated cardioids at d4.5. (E) Expression of genes representing pathways shown in Fig. 3F. Color bar: Scaled gene expression (F) Dot plots indicating GO Biological Process Term enrichment for the indicated comparisons at d4.5. (G) Complete list of pathways differentially regulated between WT, HAS2 KO and HAase-treated cardioids at d7.5. (H) Expression of genes representing pathways marked with an asterisk in (G). (I) Dot plots indicating GO Biological Process Term enrichment for the indicated comparisons at d7.5. Color bars in (A), (D) and (G): GSVA enrichment score. Colors bars in (B), (E) and (H): Scaled gene expression. Gene Ratio in (C), (F) and (I): Percentage of genes belonging to each GO term that are detected among the differentially expressed genes.

**Fig. S4.**
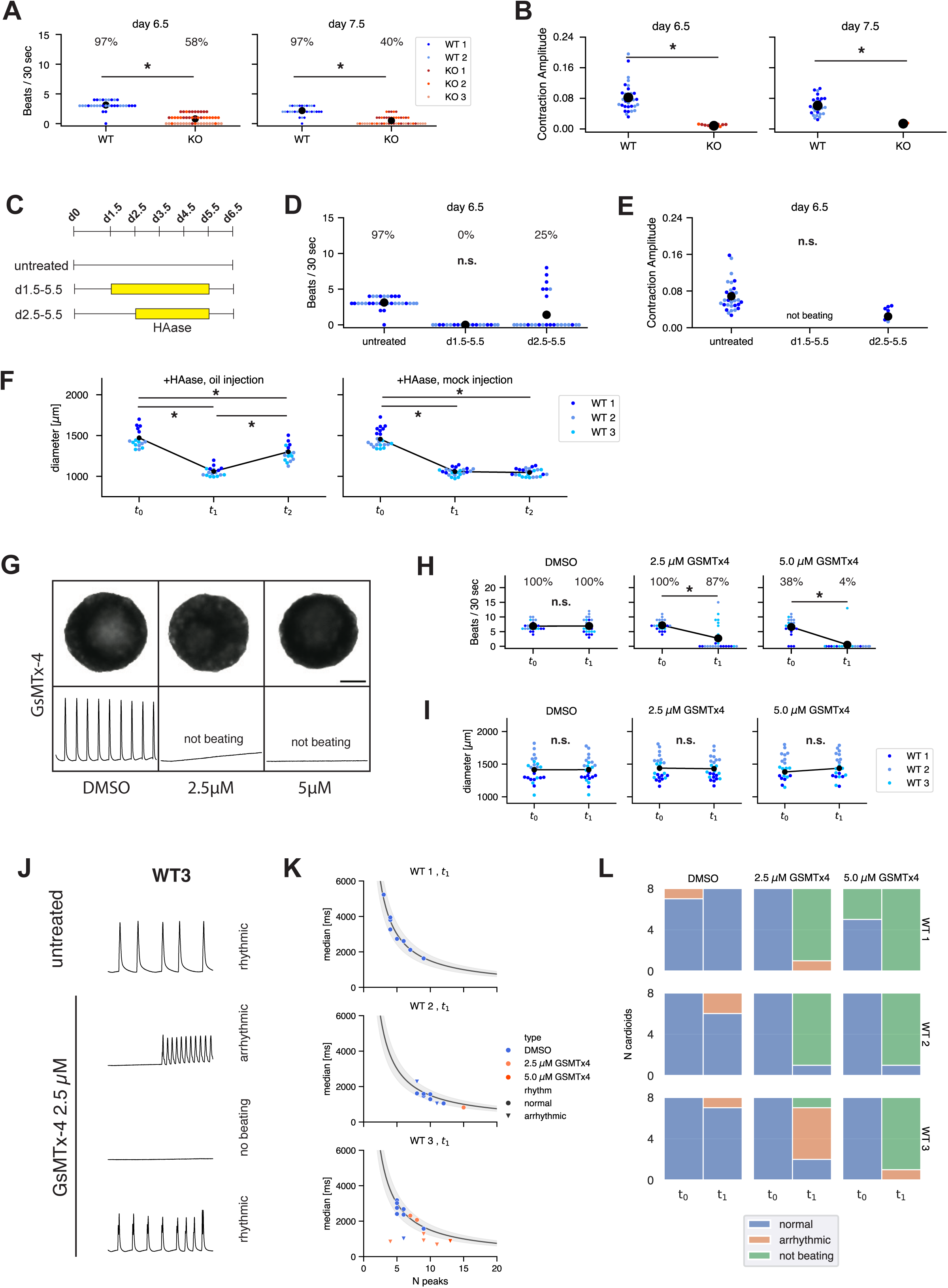
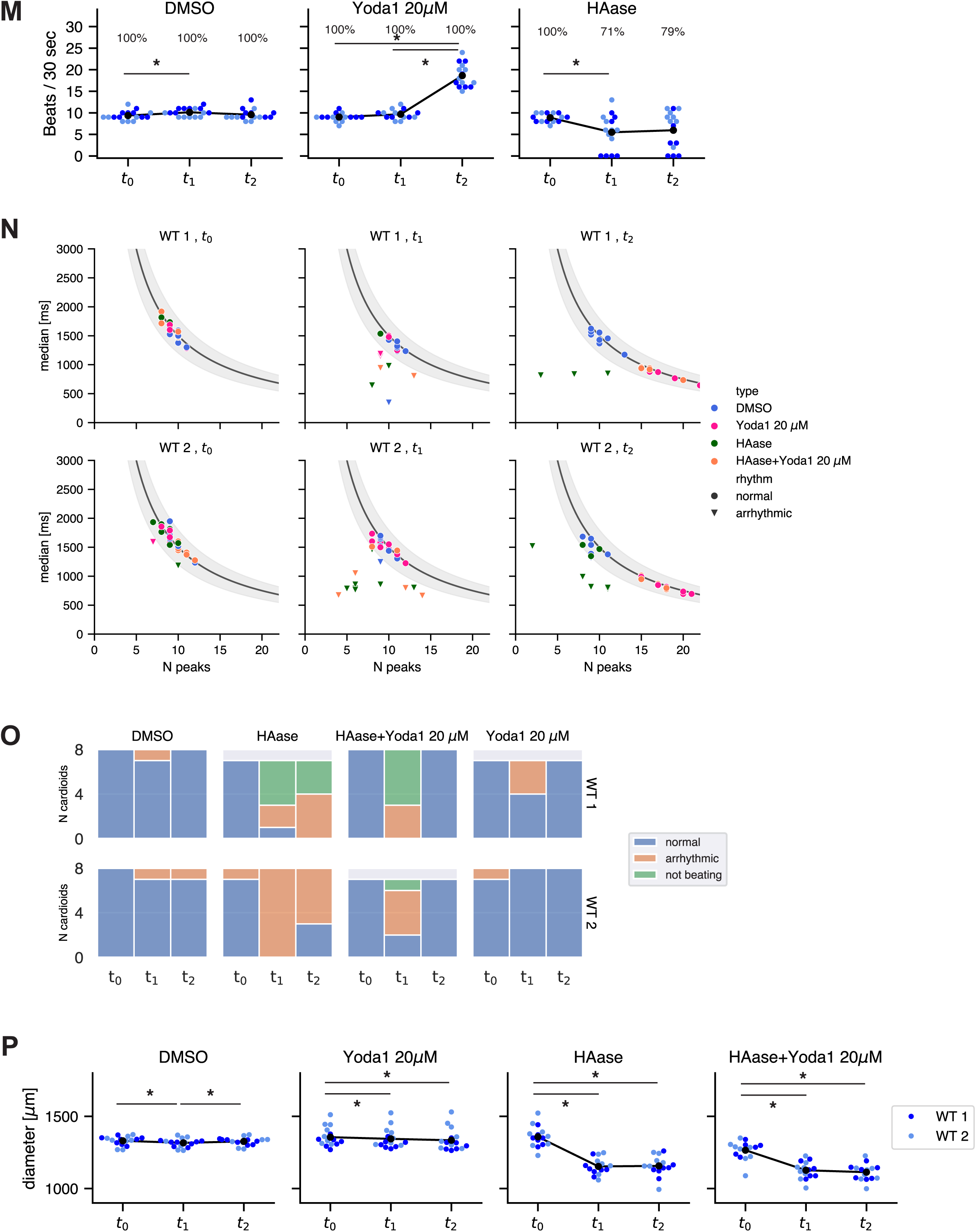
(A) Beating frequency (B) and Contraction Amplitude of WT and HAS2 KO cardioids at day 6.5 and 7.5. Same samples as Fig. 4B observed at later time points. Colors distinguish samples prepared from n=2 WT cell lines and n=3 HAS KO cell lines. n = 16 samples per cell line per condition. (C) Schematic of experiment to treat cardioids with hyaluronidase for different time windows. (D) Beating frequency and (E) Contraction Amplitude of d6.5 cardioids treated with HAase for different time windows. (F) Left: Diameter of cardioids subjected to HAase treatment, followed by oil injection. Bottom: Diameter of cardioids subjected to HAase treatment, followed by mock injection. Colors distinguish samples prepared from three cell lines. n=5-8 samples per cell line per condition. Same samples as Fig. 4E’. (G) Bright field image (top) and contraction traces (bottom) of day 5.5 cardioids treated with DMSO, 2.5 µM GsMTx4, or 5.0 µMGsMTx4. Scale bar 500 microns. (H) Beating frequency (I) and diameter of cardioids before and after GsMTx4 treatment. Colors distinguish samples prepared from three cell lines. n=7-8 samples per cell line per condition. (J) Examples of contraction traces from untreated and 2.5 µM GsMTx4 cardioids made from WT3 cell line at t_1_. The 2.5 µM GsMTx4 condition shows diversity in beating behavior, ranging from arrhythmic, no beating to rhythmic (normal) beating. (K) Comparison of the median time interval vs. number of contraction peaks N_peaks_ distinguish arrhythmic from normal beating behavior in cardioids treated with or without GsMTx4. The solid line and gray region indicates 30 sec/N_peaks_ ±20%, which is the expectation for cardioids with normal beating behavior. Samples that fall outside this range are classified as arrhythmic. Only beating samples are shown. (L) Distribution of beating behavior before and after GsMTx4 treatment. (M) Beating frequency of day 5.5 cardioids treated with either Yoda1, or hyaluronidase alone. Colors distinguish samples prepared from three cell lines. n=7-8 samples per cell line per condition. DMSO served as a carrier control for Yoda1 treatment. Same experimental cohort as Fig. 5F’. (N) Comparison of the median time interval vs. number of contraction peaks N_peaks_ distinguish arrhythmic from normal beating behavior in cardioids treated with HAase and/or Yoda1. The solid line and gray region indicates 30 sec/N_peaks_ ±20%, which is the expectation for cardioids with normal beating behavior. Samples that fall outside this range are classified as arrhythmic. Only beating samples are shown. (O) Distribution of beating behavior in cardioids treated with HAase and/or Yoda1. (P) Diameter of day 5.5 cardioids treated with either Yoda1 and hyaluronidase alone, as well as Yoda1 following hyaluronidase treatment. Colors distinguish samples prepared from three cell lines. n=7-8 samples per cell line per condition. DMSO served as a carrier control for Yoda1 treatment. Same experimental cohort as Fig. 5F’. Percentages indicate the fraction of beating samples in each condition. Asterisks indicate a statistically significant difference of p-value < 0.05.

**Fig. S5.**
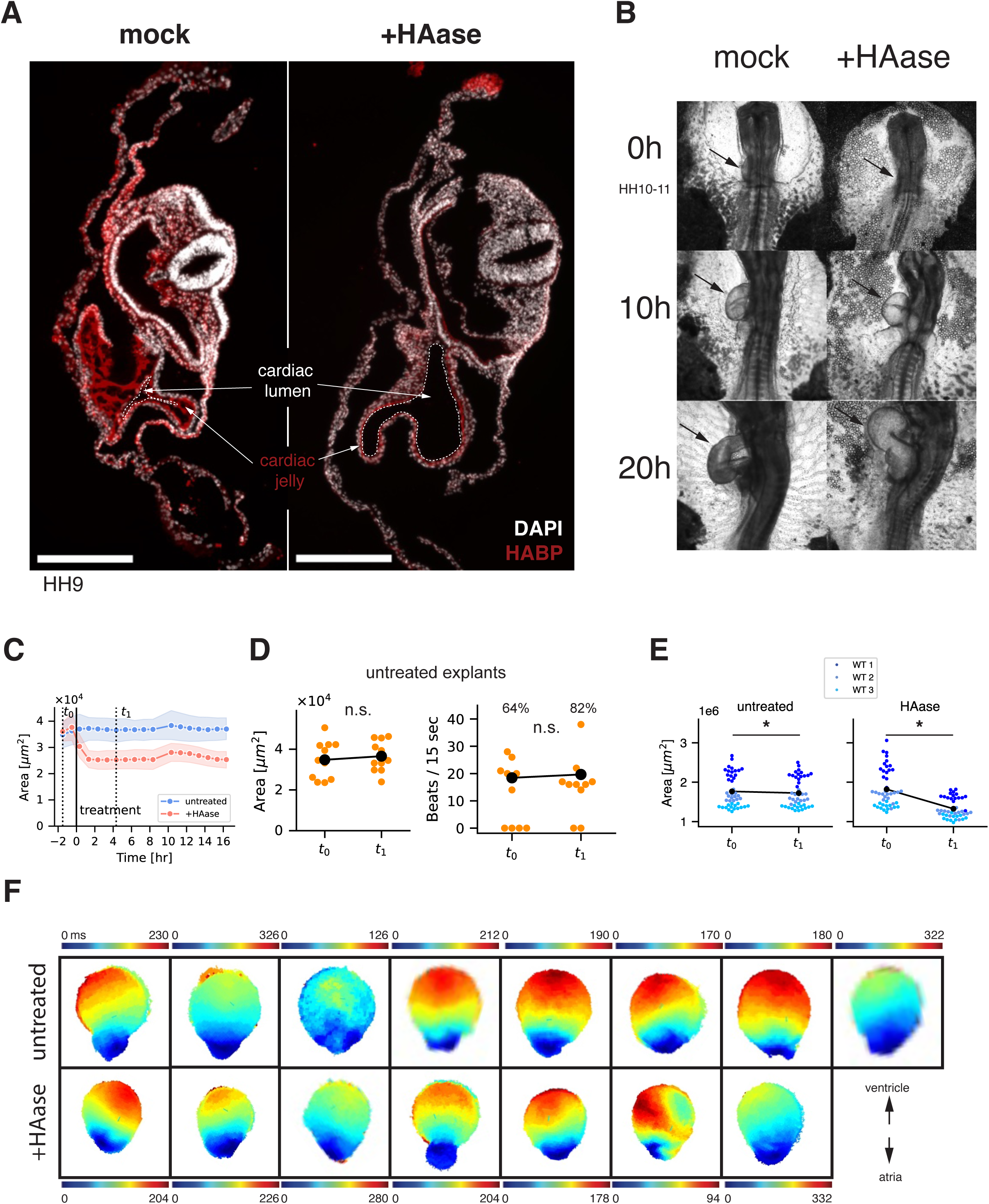
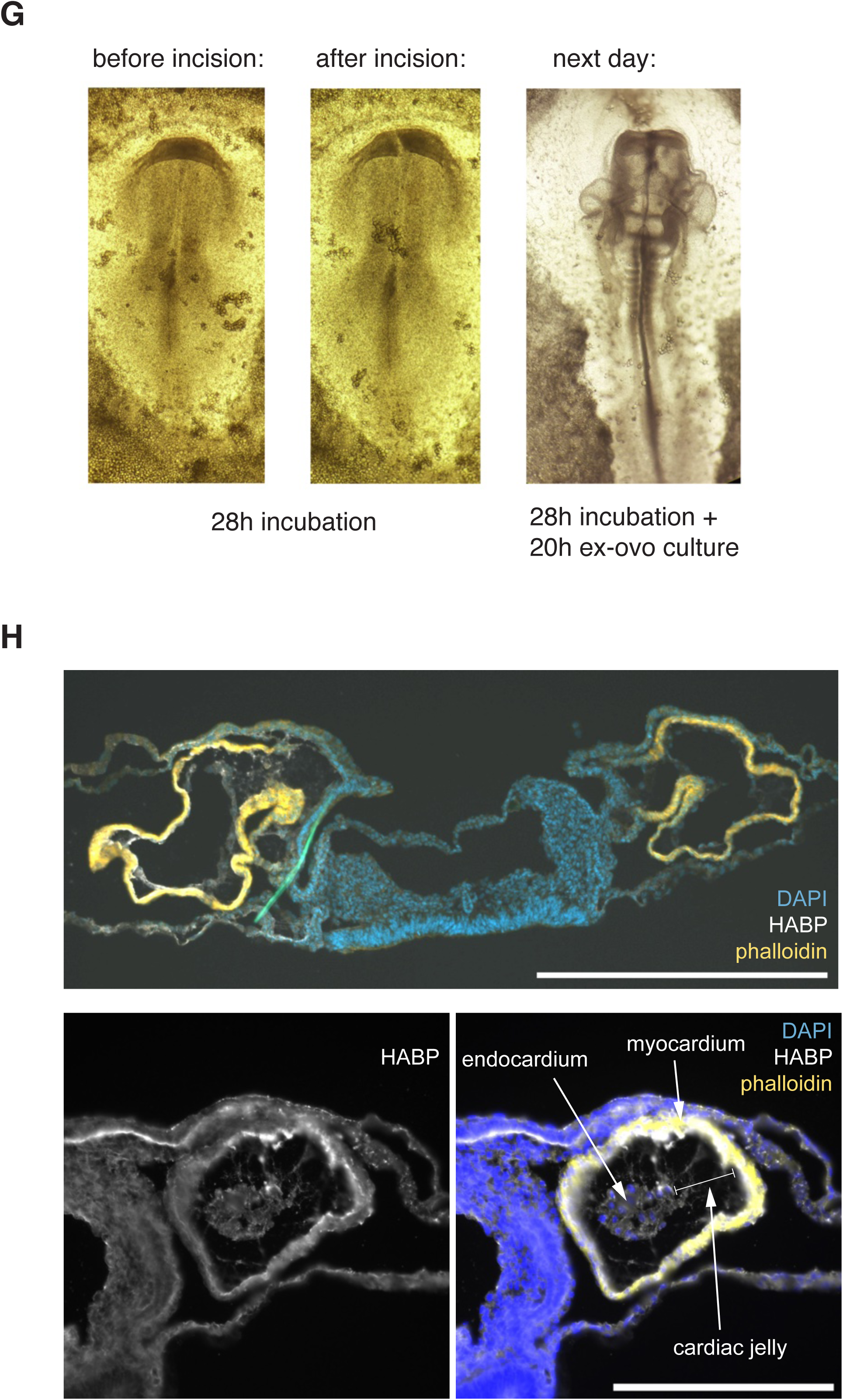
(A) Cross section through heart regions of HH9 chick embryos injected with either PBS or hyaluronidase. Note loss of cardiac jelly and expansion of the heart lumen (indicated with dashed line) in HAase-injected embryo. Scale bar 250 microns. (B) HAase injection into cardiac jelly of HH10-11 chick hearts occasionally results in drastic increase of heart volume. Timepoints shown are right after injection (0h), as well as 10h and 20h after. (C) HAase treatment leads to a reduction of tissue area of chick heart explants within two hours. Dotted lines at t_0_ and t_1_ indicate the timepoints used for statistical analysis in Fig. 5A and A’. Solid line indicates the start of treatment. (D) Control chick heart explants do not show change in tissue area or beating frequency from t_0_ to t_1_. n=11 samples. Percentages indicate the fraction of beating samples in each condition. (E) Area of LV-A multichamber cardioids before (t_0_) and after (t_1_) HAase treatment. Colors distinguish samples prepared from three cell lines. n=13-16 samples per cell line per condition. (F) Additional examples of the spatiotemporal map of calcium activation in LV-A multi-chamber cardioids showing consistent propagation from the atrial to the ventricular portion in all observed cardioids with (n=7) or without HAase (n=8) treatment. Atrial parts are oriented towards the bottom. (G) Induction of *cardia bifida* in chick embryo by foregut incision. (H) Top: Immunostaining of cryosection of *cardia bifida* chick embryo from (G) through the heart region showing a pair of phalloidin-positive heart-like structures. Scale bar 500 microns. Bottom: detail of a single heart-like structure showing HABP-positive cardiac jelly between endocardium and myocardium. Scale bar 250 microns. Asterisks indicate a statistically significant difference of p-value < 0.05.

## OTHER SUPPLEMENTARY MATERIAL

**Movie S1.** Timelapse microscopy of cavity formation, transverse (xy) plane.

**Movie S2.** Timelapse microscopy of cavity formation, sagittal (xz) plane.

**Movie S3.** Mesh visualization of cavity formation based on timelapse microscopy data.

**Movie S4.** Effect of hyaluronidase treatment between d2.5-3.5 on left ventricular cardioid size. Left: untreated, right: treated.

**Movie S5.** Effect of hyaluronidase treatment at later stages on left ventricular cardioid size. Top row: treatment between d3.5-4.5, middle row: treatment between d4.5-5.5, bottom row: treatment between d5.5-6.5. Left: untreated, right: treated.

**Movie S6.** Effect of HAS2 KO on left ventricular cardioid beating at d5.5. Note very weak twitching motions in KO1 and KO2. Video sped up 2x.

**Movie S7.** Effect of long-term hyaluronidase treatment on left ventricular cardioid beating at d5.5. Left: untreated, middle: hyaluronidase treatment from d1.5-5.5, right: hyaluronidase treatment from d2.5-5.5. Note very weak twitching motions in sample treated with HAase from d2.5-5.5. Video sped up 2x.

**Movie S8.** Acute effect of hyaluronidase treatment and oil injection on left ventricular cardioid beating at d5.5. Left: before hyaluronidase treatment, middle: after hyaluronidase treatment, right: after hyaluronidase treatment and oil injection. Video sped up 2x.

**Movie S9.** Effect of GsMTx-4 on left ventricular cardioid beating at d5.5 Video sped up 2x.

**Movie S10.** Acute effect of hyaluronidase treatment and Yoda1 on left ventricular cardioid beating at d5.5. Video sped up 2x.

**Movie S11.** Effect of acute hyaluronidase treatment on beating of chick heart explants. Top: untreated, bottom: HAase-treated. t_0_: 1.5h before medium change, t_1_: 4.3 hours after medium change.

**Movie S12.** Effect of acute hyaluronidase treatment on beating of fused left ventricular-atrial cardioids at d7.5. Left: before hyaluronidase, right: after hyaluronidase. Video sped up 2x.

**Movie S13.** Effect of acute hyaluronidase treatment on calcium transients of cardioids top row: left ventricular cardioid, middle row: atrial cardioid, bottom row: fused left ventricular-atrial cardioid. left: before hyaluronidase, right: after hyaluronidase. Video sped up 2x.

**Table S1.** Bulk RNA-seq differential expression results for all pairwise comparisons across WT, HAS2 KO, and HAase-treated LV cardioids.

**Table S2.** Gene Ontology Biological Process enrichment analysis for differentially expressed genes from the indicated transcriptomic comparisons.

### Cell lines used in experiments

| Figure and panel | Cell line |
| --- | --- |
| Fig. 1 B | WT1: WTC WT<br>WT2: WTC TNNT2-GFP<br>WT3: WTC WT |
| Fig. 1 C, C', D, D' | WTC CAG-GFP |
| Fig. 1 E | WTC WT |
| Fig. 1 F | WTC WT |
| Fig. 1 H | WTC CTNNB1-GFP |
| Fig. 2 A, A' | WTC TNNT2-GFP |
| Fig. 2 B | WTC WT |
| Fig. 2 D, D' | WT1: WTC TNNI1-GFP<br>WT2: WTC MYL7-GFP |
| Fig. 2 E | WT: WTC WT<br>KO: WTC HAS2 KO 1 clone 4 |
| Fig. 2 F, F' | WT: WTC WT |
| Fig. 3 H | WTC TNNI1-GFP |
| Fig. 3 I | WTC TNNI1-GFP |
| Fig. 4 B', B'', fig. S4A, B | WT1: WTC TNNI1-GFP<br>WT2: WTC MYL7-GFP |
| Fig. 4 C', C'', fig. S4D, E | WT1: WTC TNNI1-GFP<br>WT2: WTC CDH5-Tomato/MYL7-GFP |
| Fig. 4 E', fig S4F | WT1: WTC TNNI1-GFP<br>WT2: WTC CDH5-Tomato/MYL7-GFP<br>WT3: WTC FUCCI |
| fig. S4H, I, J, K, L | WT1: WTC TTN-GFP<br>WT2: WTC TNNI1-GFP<br>WT3: WTC TNNT2::GCaMP |
| Fig. 4 F', F'', fig. S4M, N, O, P | WT1: WTC TNNI1-GFP<br>WT2: WTC TNNT2::GCaMP |
| Fig. 5 B', B'', D', fig. S5D, E | WT1: WTC TNNI1-GFP<br>WT2: WTC CAG-GFP<br>WT3: WTC TNNT2::GCaMP |

## MATERIAL AND METHODS

### Human pluripotent stem cells (hPSCs)

All used cell lines are derived from the WTC iPS cell line (male, skin fibroblast-derived), which was developed at Dr. Bruce R. Conklin’s laboratory (Gladstone Institute of Cardiovascular Disease, UCSF, USA) and purchased from the Coriell Institute for Medical Research (USA). The Allen Institute for Cell Science’s cell lines are derived from the WTC11 cell line and received from the Coriell Institute for Medical Research (USA).

### Cell lines used in this study

| Cell line | Source | Identifier |
| --- | --- | --- |
| Wild-Type WTC iPSC | Coriell Institute for Medical Research | GM25256 |
| MYL7-GFP WTC iPSC | Allen Institute for Cell Science | AICS-0052-003 |
| TNNI1-GFP WTC iPSC | Allen Institute for Cell Science | AICS-0037-172 |
| TTN-GFP WTC iPSC | Allen Institute for Cell Science | AICS-0048-039 |
| CTNNB1-GFP WTC iPSC | Allen Institute for Cell Science | AICS-0058-067 |
| TNNT2-GFP WTC iPSC | HeartBeat.Bio | in-house |
| TNNT2::GCaMP6f WTC iPSC | HeartBeat.Bio | in-house |
| WTC CAG-GFP | this paper | N/A |
| WTC FUCCI | this paper | N/A |
| WTC MYL7-GFP/CDH5-Tomato | Hofbauer et al, 2021 | N/A |
| WTC HAS2 KO 1 clone 4 | this paper | N/A |
| WTC HAS2 KO 2 clone 14 | this paper | N/A |
| WTC HAS2 KO 3 clone 51 | this paper | N/A |
| HAS2KO; TetOn-GFP-HAS2<br>WTC iPSC | this paper | N/A |

### hPSC culture

Cells were maintained in E8 culture system(*57*) containing 0.5% BSA (Europa Biosciences #EQBAH70), thermostable FGF2-G3 (5.5–10 ng/mL; Qk053), and TGFβ1 (1.8 ng/mL; R&D #240-B-010). Cells were cultured on Vitronectin XF-coated TPP plates (#92012). Passaged every 2–4 days at ∼70% confluence using 4 minutes incubation in TrypLE Express (Gibco #12605010). ROCKi (Y-27632, 5 µM; Tocris #1254) was added to the medium during the first day after splitting. Cells were kept in a humidified incubator at 37°C, 5% CO2 and normoxia conditions.

### Generation of cardioids

To generate cardioids, hPSCs at ∼70% confluency were seeded at 25-35k cells per well (24-well plates, TPP #92024) in E8 + ROCKi (5 µM). 24 hours after seeding (d0) cells were induced using mesoderm induction media. 36-40 hours later (d1.5) cells were dissociated using 2 minutes incubation in TrypLE Express and reseeded into ultra-low attachment 96-well plates (Corning #7007) at 15-20k cells/well in Cardiac Mesoderm Patterning Media 1 supplemented with ROCKi (5 µM). After seeding, cells were spun down at 200g for 4 minutes. 24 hours later (d2.5) media was replaced with Cardiac Mesoderm Patterning Media 1 without ROCKi. The following two days (d3.5-d4.5) medium was changed daily to Cardiac Mesoderm Patterning Media 2. For the subsequent two days medium was changed daily to Cardiomyocyte Differentiation Media. All media formulations are described below.

### Cardioid compartment-specific differentiation media

All differentiation media are based on chemically defined medium (CDM) consisting of 5 mg/ml bovine serum albumin (Europa BiosciencesEQBAH70) in 50% IMDM (Gibco #21980065) plus 50% Ham’s F12 Nutrient-MIX (Gibco #31765068), supplemented with 1% concentrated Lipids (Gibco #11905031) 0.004% Monothioglycerol (Sigma #M6145-100ML) and 15 mg/ml of Transferrin (Roche #10652202001), 1X Antibiotic-Antimycotic (Thermo Scientific #15240062).

Mesoderm Induction Media (d0-d1.5)

*Left ventricular cardioid (LV):* CDM containing FGF2-G3 (5.5 ng/mL, Qk053), LY294002 (5 µM, Tocris #1130), Activin A (5ng/ml, Cambridge University), BMP4 (10 ng/ml, R&D Systems RD-314-BP-050), and CHIR99021 (3 µM, R&D Systems RD-4423/50)

*Atrial cardioid (A):* CDM containing FGF2-G3 (5.5 ng/mL), LY294002 (5 µM), Activin A (50 ng/ml), BMP4 (10 ng/ml), and CHIR99021 (4 µM)

Cardiac Mesoderm Patterning Media 1 (d1.5-d3.5)

*LV:* CDM containing BMP4 (10 ng/ml), FGF2-G3 (1.466 ng/mL), Insulin (10 µg/ml), C59 (2 µM, Tocris #5148/10) and Retinoic Acid (50 nM, Sigma Aldrich #R2625).

*A:* CDM containing SB 431542 (10 µM, Tocris #1614), XAV939 (5 µM, SelleckChem #S1180) and Retinoic Acid (500 nM).

Cardiac Mesoderm Patterning Media 2 (d3.5-d5.5)

*LV:* CDM containing BMP4 (10 ng/ml), FGF2-G3 (1.466 ng/mL), Insulin (10 µg/ml), C59 (2 µM) and Retinoic Acid (50 nM).

*A:* CDM containing BMP4 (10 ng/ml), FGF2-G3 (1.466 ng/mL), Insulin (10 µg/ml), XAV939 (5 µM) and Retinoic Acid (500 nM).

Co-development Patterning Media (d3.5-d5.5)

CDM containing BMP4 (10 ng/ml), FGF2-G3 (1.466 ng/mL), Insulin (10 µg/ml), C59 (2 µM) and Retinoic Acid (500 nM).

Cardiomyocyte Differentiation Media (d5.5-d7.5)

*LV&A:* CDM containing BMP4 (10 ng/ml), FGF2-G3 (1.466 ng/mL) and Insulin (10 µg/ml)

### Generation of multi-chamber cardioids

Atrial and left ventricular cardioids were differentiated according to their specific protocols. To visually distinguish atrial from left ventricular portions in multi-chamber cardioids, atrial cardioids were incubated at d2.5 in medium supplemented with DiIC18(5)-DS (0.5 µg/ml, ThermoScientific #D12730) for one hour, followed by two washes in the same medium. At d3.5 atrial and ventricular cardioid fusion was accomplished by combining them in a single well of ultra-low attachment 96-well plates. Medium was subsequently changed to *Co-development Patterning Media* and replaced again at d4.5. Between d5.5-d7.5 medium was changed to *Cardiomyocyte Differentiation Media*.

### Generation of CAG-GFP cell line

The CAG-GFP expression cassette was integrated into the AAVS1 locus via TALENs using pAAV-Puro_CAG-EGFP (a gift from Alessandro Bertero). hPSCs that were approximately 70% confluent, were dissociated using TrypLE Express and transfected using the P3 Primary Cell 4D-Nucleofector X Kit S (Lonza-BioResearch #V4XP-3032) and Amaxa 4D-Nucleofector (Lonza-BioResearch) with pulse code DS-120. Post nucleofection, cells were incubated in E8 supplemented with ROCKi (5mM) on a 6-well plate previously coated with Vitronectin XF (StemCell Technologies #7180). The following day, medium was refreshed with E8 supplemented with ROCKi. The media was replaced with E8 for the following two days. Antibiotic selection was carried out for three days with daily media changes using E8 supplemented with Puromycin (0.5µg/ml). After selection, cells were dissociated using TrypLE Express and seeded as single cells (2k cells) into a 10cm dish. Colonies were picked, transferred to a 96 well plate and GFP signal was confirmed using a widefield microscope. Using TrypLE Express selected colonies were dissociated and expanded to a 12 well plate for use in experiments.

### Generation of HAS2 KO

HAS2 was knocked out in WTC WT cells using CRISPR/Cas9 multi-guide approach (Synthego KO kit). Guide sequences: sgRNA0: CAAACAGGCUUUGGAUGAUG, sgRNA1: AAUUAUCCGUUUGGAUAAAC, sgRNA2: GUGUGGUUCCAAUUAUUCUC. Once hPSCs were approximately 70% confluent they were dissociated using TrypLE Express and transfected using the P3 Primary Cell 4D-Nucleofector X Kit S (Lonza-BioResearch # V4XP-3032) and Amaxa 4D-Nucleofector (Lonza-BioResearch) with pulse code DS-120. Post nucleofection, cells were incubated in E8 supplemented with ROCKi (5mM) on a 6-well plate previously coated with Vitronectin XF (StemCell Technologies #7180). The following day, medium was replaced with E8 (supplemented with ROCKi the first day after transfection). Once cells were approximately 70% confluent, cells were dissociated using TrypLE Express and seeded as single cells (4k cells) on a 10cm dish. Single colonies were split into duplicate 96w plates and genotyped using gel electrophoresis and inhouse Sanger sequencing. Gene editing was analyzed using Synthego’s online tool ICE (https://ice.synthego.com/<u>#/</u>). From the initial screening three independent HAS2 KO clones were expanded and their genotype confirmed a second time before using them in subsequent experiments.

### Generation of Tet-On HAS2-GFP and HAS2-GFP overexpression

As a parental line we targeted HAS2 KO1 (fig. S2B) line. Human HAS2 ORF (CCSB Human ORFeome collection Clone ID: 13942) was PCR amplified to produce a N-terminal GFP fusion vector pME-GFP-HAS2. Gateway LR Clonase (Invitrogen) was used to insert GFP-HAS2 fusion protein into the tetOn destination vector PiggyBac (PB)-tetOn-destination-PGK-hygro(*58*). The resulting PiggyBac (PB)-tetOn-GFP-HAS2-PGK-hygro and PB-CAG-rtTA3-PGK-puro vector(*58*) integrated into the genome of HAS2 KO human iPSCs assisted by the non-integrating PBase transposase vector. All three plasmids were delivered using P3 Primary Cell 4D-Nucleofector X Kit S (Lonza-BioResearch #V4XP-3032) and Amaxa 4D-Nucleofector (Lonza-BioResearch). After transfections, we performed selection with 0.5 µg/ml Puromycin and 50 µg/ml Hygromycin B and picked clones to establish cell lines. GFP expression upon doxycycline treatment was validation of cell lines. To induce expression of HAS2-GFP, doxycycline (1 µg/ml, Sigma-Aldrich #D9891) was added to the culture media from d1.5-5.5.

### Hyaluronan molecular weight analysis by single-molecule solid-state nanopore

Left ventricular cardioids collected at day 6.5 were rinsed three times in PBS and the supernatant was fully removed. Pellets were resuspended in 0.4 mL Proteinase K solution (Invitrogen cat. 433793) and incubated overnight at 37°C with gentle agitation. Samples were stored at 4 oC until further processing. Cleanascite LX (Biotech Support Group, Cat. LX155-10) was added to the sample at a 1:1 ratio, mixed for 20 min, and centrifuged at 5000 × g for 10 min to remove lipids and cell debris and the supernatant was recovered. This was subjected to phenol:chloroform extraction (25:24:1; Fisher Scientific, Cat. AC327111000) to isolate HA, glycans, and nucleic acids in the aqueous phase. The same procedure was repeated twice with pure chloroform (Cat. 60-047-880) to remove remnant phenol. Affinity extraction and nanopore analysis were conducted following protocols reported elsewhere(*59*, *60*). Briefly, samples were incubated for 60 min at room temperature with 1.5 mg magnetic beads (Dynabeads™ M-280 Streptavidin, Thermo Fisher, Cat. 11205D) conjugated with biotinylated Versican G1 domain (Echelon Biosciences, Cat. G-HA02). The beads were pulled down magnetically, washed with 1× PBS, and bound HA was eluted using 40 µL measurement buffer (6 M LiCl, 10 mM Tris, 1 mM EDTA, pH 8.0) for 60 min at room temperature. The eluted HA was analyzed using a 30 nm-thick, prefabricated solid-state nanopore (12.2 nm diameter, Norcada, P/N: NXPR4002X-16nm-ABX1) under a 300 mV applied bias. Events were detected by in-house software (Labview) using a threshold of 5X the RMS baseline noise. Individual translocation event areas (event charge deficit, or ECD) were converted to molecular weight through an internal calibration measurement of three quasi-monodisperse HA standards(*61*) (111, 545, and 1071 kDa; Hyalose, LLC). Only events between 50 kDa and 10 MDa were included in the analysis.

### Enzymatic perturbation using hyaluronidase

For enzymatic perturbation cardioids were treated with hyaluronidase (1 mg/ml in the corresponding media, Sigma-Aldrich #H3506) at 37C, 5% CO2. Long-term treatments (Fig. 4C, C’, C’’) started on either d1.5 or d2.5 and lasted until d5.5. Acute treatments were done at d5.5 for left ventricular cardioids and at d7.5 for atrial and LV-A cardioids an hour before assaying contractility.

### Mechanical rescue by oil-droplet injection

Fluorinated oil (HFE-7500, fluorochem F051243) was injected into HAase-treated cardioids approximately 2-3 hours after treatment. Microinjection was manually done via mouth-pipette using needles (TW100F-4, World pPrecision Instruments) pulled on a Micropipette Puller (Sutter P-97) (heat 520/pull 150/vel 100/time 150). The final volume of injected oil was adjusted to approximate the control (untreated) cardioid size. Cardioids that did not retain their oil droplet because of leakage/rupturing of the tissue were excluded from further analysis. Mock injection was performed by puncturing cardioids with a needle.

### Pharmacological modulation of stretch-sensitive channels

Pharmacological inhibition of stretch-activated channels was performed using GsMTx4 (MedChemExpress #HY-P1410). LV cardioids were treated with 2.5 µM or 5 µM GsMTx4. Treatment was initiated 1 hour before the start of contraction recordings. Because GsMTx4 was diluted in DMSO, parallel controls were included using media with and without BSA, as well as a DMSO-only control. BSA was excluded from the treatment media. Pharmacological activation of Piezo1 was induced using Yoda1 (Tocris #5586/10). Cardioids were treated with 20 µM Yoda1 on day 5.5 (LV), or day 7.5 (LV-A). Treatments were started 2 hours before initiating contraction assays.

### Contractility and arrhythmia analysis of cardioids

1-2 hours before recordings media were refreshed and the 96-well plate was transferred to an environmentally controlled stage incubator (37C, 5% CO2, water-saturated air atmosphere, Okolab Inc, Burlingame, CA, USA) and imaged using a widefield microscope (Axioobserver Z1 (inverted) with sCMOS camera). Each well was imaged at either 50 or 100 frames per second for a duration of 30 seconds. Videos were then analyzed using MUSCLEMOTION (*62*); the data was read into custom-made software for reported calculations. We manually recorded samples that only showed one contraction peak. The contraction amplitude is the amplitude given from MUSCLEMOTION divided by the size of the cardioid.

Our criteria for arrhythmia was based on the number of contractile peaks recorded during a 30 sec recording window (N_peaks_) and the median time interval between peaks (T_interval_); if T_interval_ was outside the range of 30 sec/N_peaks_ ± 20%, the beating pattern was classified as “arrhythmic”. If T_interval_ lied within this range, the beating pattern was classified as “normal”. Cardioids with N_peaks_=0 were classified as “not beating”.

### Transcriptomic profiling (Bulk RNA-seq)

A total of 8 cardioids were pooled per replicate and condition. For each condition cardioids from at least 3 biological replicates were included. RNA was extracted using an in-house RNA bead isolation kit semi-automated using KingFisher devices (KingFisher Duo Prime). Libraries were prepared according to the manufacturer’s instruction using QuantSeq 3‘ mRNA-Seq Library Prep Kit for Illumina (FWD) (Lexogen #015). Samples were checked for adequate size distribution with a fragment analyzer (Advanced Analytical Technologies, Inc). The RNA-seq library was submitted to the Vienna Biocenter Core Facilities (VBCF) Next-Generation-Sequencing (NGS) facility for sequencing (NovaSeq SP SR100 XP).

### Differential gene expression analysis and functional enrichment

Initial R1 read processing was performed using BBDuk v38.06 by removing adapter, polyA and low-quality bases from the 3’ end (ref=polyA.fa.gz,truseq.fa.gz k=13 ktrim=r useshortkmers=t mink=5 qtrim=r trimq=10 minlength=20) and filtering for abundant sequences included in the iGenomes UCSC hg38 reference (human rDNA, human mitochondrial chromosome, phiX174 genome, adapter) (k=31). The processed R1 reads were analyzed using genome and gene annotation for the GRCh38 assembly obtained from *Homo sapiens* Ensembl release 94. Reads were aligned to the genome using star v2.6.0c and reads in genes were counted with featureCounts (subread v1.6.2). Differential gene expression analysis on raw counts and variance-stabilized transformation of count data for heatmap visualization were performed using DESeq2(*63*) v1.18.1.

### Functional annotation of differentially expressed genes

Differentially expressed genes were selected based on p-adj < 0.05. The clusterProfiler R package (*64*) v4.12.0 was used to perform Gene Ontology (GO) enrichment analysis using the enrichGO() function. Redundant GO terms were removed using the simplify() function.

Enriched GO terms (adjusted p-value < 0.05) were retained only if supported by at least three genes from the input gene list and if these genes accounted for ≥2% of all genes annotated to the respective GO term. The Odds Ratio was calculated for each GO term based on the formula: Odds Ratio = (a/b) / (c/d) = a*d / b*c, where a: number of differentially expressed genes belonging in the GO term, b: number of differentially expressed genes not belonging in the GO term, c: GO term size, d: number of all background genes – GO term size. The top 30 GO terms, based on Odds Ratio, were eventually selected for each comparison. Activity of selected enriched GO terms was assessed per sample using the GSVA R package (*65*) v1.52.3. Only genes from the input gene list that contributed to each respective GO term were used in this analysis.

### Fixation and cryosectioning

Cardioids were fixed for 30 minutes at room temperature with 4% PFA in PBS and subsequently washed 3×10 minutes in PBS. For cryoprotection, cardioids were infiltrated with 30% sucrose in PBS overnight at 4C, or until they sank to the bottom of the tube. Samples were then embedded in cryomolds filled O.C.T. medium (Scigen #4586) and frozen by placing the mold on a metal block, which has been pre-cooled in liquid nitrogen. 14um sections were made on a cryostat (Epredia Cryostar NX70), collected on slides (Leica Permaflex Plus) and kept at-20C until further processing. Chick embryos were fixed with 4% PFA in PBS for 2.5 hours on ice and washed 3×30mins in PBS. Fixed tissues were cut into smaller pieces and processed for cryosectioning in the same way as cardioid samples.

### Immunostaining and imaging of cryosections

Slides were equilibrated at room temperature for 10 minutes and washed in PBS for 15 minutes. Samples were then blocked in blocking solution (4% donkey (Bio-Rad Laboratories C06SC) or goat serum (Bio-Rad Laboratories C07SA in PBS 0.1% Triton X-100 for 30 minutes. Slides were then incubated in primary antibody (in blocking buffer) overnight at 4C. On the following day samples were washed 3×10 minutes in PBS 0.1% Tween 20, followed by incubation in secondary antibody (in blocking buffer) for 1 hour at room temperature protected from light. Finally, slides were washed 3×10 minutes in PBS 0.1% Tween, mounted in fluorescent mounting medium (Dako Agilent Pathology Solutions #S3023), dried overnight at room temperature, protected from light and stored at 4C until imaging. Stained slides were either imaged using Slidescanner (Pannoramic FLASH 250 III Scanner) or upright widefield microscope (Axio Imager.Z2 with sCMOS camera).

### List of antibodies and other proteins used for immunostaining

| **Antibody/protein** | **Source** | **Identifier** |
| --- | --- | --- |
| DAPI | Sigma-Aldrich | #D9542 |
| HABP-Biotin | Merck | #385911 |
| Streptavidin Alexa 647 | Invitrogen | #S32357 |
| NKX2-5 | R&D Systems | #AF2444 |
| Troponin T | Thermo Scientific | #MS-295-P |
| Donkey anti-Mouse Alexa 594 | Thermo Scientific | #A21203 |
| Donkey anti-Goat Alexa 647 | Thermo Scientific | #A21447 |
| Phalloidin | Invitrogen | #A12380 |

### Wholemount 3D imaging of cardidoids

For 3D wholemount imaging, cardioids were fixed in 4% PFA and labelled with the pan-protein dye AlexaFluor 647 NHS ester (Thermo A37573) at 20 µg/ml in 0.1M sodium phosphate buffer pH 8.0. After washing samples in PBS to remove excess dye, samples were optically cleared with CUBIC R2 without TEA (*66*, *67*). Cleared samples were imaged with the Yokogawa CQ1 microscope equipped with a 10X 0.45 NA air objective lens, Yokogawa CSU-W1 spinning disc confocal head, Hamamatsu Flash 4.0 sCMOS camera, 640 nm excitation light, and 708/75 nm bandpass emission filter. Z-stacks were acquired with 2 micron z-steps and up to 551 planes. For larger samples, we performed 2×2 tiled imaging.

### 3D morphological analysis of cleared cardioids

To quantify 3D morphology (*68*), namely cavity and cardioid volume in optically cleared samples, z-stacks were converted to isotropic images of voxel size 2.90 microns using the makeIsotropic function in CLIJ2(*69*) in Fiji (*70*), assuming an original pixel size of 0.645 micron in xy and 2 x 1.45 = 2.90 micron in z, the latter correcting for the refractive index mismatch between air and CUBIC R2. Using the isotropic images, we segmented cell-containing and non-containing regions with the pixel classification workflow in Ilastik (*71*) and exported the probability image of the cell containing region. The subsequent steps were all performed with custom Python scripts using scikit-image (*72*): The 3D tissue probability maps were smoothed with a Gaussian filter and binarized using Otsu’s threshold to generate tissue masks. Small isolated objects (volume < 5.8 × 10⁵ µm³) were removed to exclude debris or small clusters of floating cells. In some samples, we observed acellular spaces within the tissue mask that were connected to the surrounding. To fill these open spaces, we processed the tissue masks with the scikit-image fill holes function in a plane-by-plane manner along the z-axis to generate binary cardioid masks. The binary difference between the filled cardioid and the tissue masks defined the cavity mask. We converted the 3D binary masks (tissue, cardioid, cavity) into 3D labelled images and converted each object into surface meshes using the marching cubes algorithm. For each mesh object, we computed the volume, surface area, and Euler characteristic using functions available in trimesh. PyVista (*73*) was used for mesh visualization.

### Live imaging of cavity formation

CAG-GFP WTC iPSCs were used to generate left ventricular cardioids using an alternative direct 3D method (*14*). Specifically, 2.5k cells/well were directly seeded into mesoderm induction media supplemented with ROCKi (5 µM) in ultra-low attachment 96-well plates (Corning #7007). 21 hours after seeding, cardioids were transferred to custom imaging chambers and subjected to live imaging on a light sheet microscope (Viventis LS1) with a 25X 1.1NA water immersion objective lens, 0.75x demagnifying tube lens, and 488 nm excitation laser. Every 20 min, *z* stacks were acquired at 3 µm intervals, covering a sample depth of 600 µm. 40 hours after seeding, the medium was replaced with Cardiac Mesoderm Patterning Media 1.

### 3D morphological analysis of lightsheet microscopy data

To quantify cavity and cardioid volume in lightsheet microscopy data, z-stacks were converted to isotropic images of voxel size 3.00 microns as described above, assuming an original pixel size of 0.347 micron in xy and 3.00 micron in z. Using the isotropic images, cell-containing regions were segmented using a custom Python script. For each time point, images were first smoothed using a low-pass Butterworth filter to reduce high-frequency noise. Segmentation thresholds were manually selected at every fifth time point and linearly interpolated for the remaining time points, resulting in a binary tissue mask. We followed the same steps for removing small objects, defining cavities, and volume quantification as described above.

### Scanning electron microscopy

Cardioids were transferred to a mixture of 2% Paraformaldehyde (Electron Microscopy Sciences, Hatfield, PA) and 2% Glutaraldehyde (Agar Scientific, Essex, UK) in 1x PBS and incubated overnight at room temperature. Organoids were cut into halves to obtain images from the inner regions. All organoids were post-fixed in 2% osmium tetroxide (Agar Scientific, Essex, UK) for 40 min at 4°C. Samples were then 2x rinsed with PBS and 1x in ddH2O (10 min each). Chemical dehydration was performed in a 3:1 mixture of 2,2-Dimethoxypropane (DMP, Sigma-Aldrich) and ddH2O for 10 min, followed by three times anhydrous acetone and three times a 1:1 mixture of anhydrous acetone and Hexamethyldisilazane (HMDS, Sigma-Aldrich), 30 min each step. After two additional dehydration steps in pure HMDS for 1h each, samples were air dried, sputtered with a thin gold layer in a Balzers SCD 050 and examined with a Hitachi TM-1000 tabletop scanning electron microscope operated at 17 kV and equipped with a high-sensitive semiconductor BSE detector.

### Dextran penetration assay

Cardioids were incubated in *Cardiac Mesoderm Patterning Media 2* supplemented with FITC Dextran 150 kDa (FD150S Sigma-Aldrich) at d5 and imaged with a lightsheet microscope (Viventis LS1) at d6 using the 488 nm excitation laser.

### Cardioid diameter measurements

To quantify the diameter of cardioids from bright field images, we used a CellProfiler(*74*) pipeline that incorporated the modules ImageMath (inversion), IdentifyPrimaryObjects, and MeasureObjectSizeShape.

### Widefield imaging of live cardioid samples

Live cardioids in 96-well plates were either imaged using Celigo Imaging Cytometer microscope (Nexcelom Bioscience, LLC), Axioobserver Z1 (inverted) with sCMOS camera, or CellDiscoverer 7 widefield microscope.

### Analysis of Ca^2+^ signal

To follow Ca^2+^ propagation across LV–A fusions during cardioid contraction, a WTC line expressing GCaMP6f (WTC TNNT2::GCaMP) was used (*15*). Calcium imaging data were processed using the AQuA2 analysis platform (*75*). For each cardioid, a substack of 50 frames was extracted from the original recording. Rising maps were generated from this substack to identify the onset of Ca²^+^ signals. The timing of signal propagation across chambers was calculated by multiplying the frame number corresponding to the rising phase by the exposure time (20 ms per frame).

### Chick embryo assays

All chicken embryo experiments were conducted complying with Austrian and European legislation. Wild-type fertilized eggs (*Gallus gallus*) were obtained from Schropper GmbH (Gloggnitz, Austria) and stored at 18C for a maximum of one week, before incubating them in a humidified chamber at 37C.

For heart explantation, eggs were opened after 48 hours of incubation and the embryos were staged, sectioned and transferred to a dish containing PBS. Explanted hearts were immediately transferred to a 96w plate filled with 150 µl of CDM supplemented with Antibiotic-Antimycotic (1X). After that heart explants were transferred to the incubator (37C, humidified, 5% CO2) of the microscope and equilibrated for 2 hours. After the first recording for the baseline contractility, either 50 µl of CDM or CDM supplemented with hyaluronidase (final concentration 1 mg/ml) was added to the explants. The second recording took place approximately 4.5 hours after hyaluronidase treatment. All brightfield microscopy images were recorded at 69 frames per second (fps) using a Zeiss Cell Discoverer equipped with a 5x objective lens. Images were converted to MP4 video files of 23 fps and analyzed with BeatProfiler (*76*) with settings (invert: True, method: mean fft, limit memory: False) to detect contraction peaks. To measure the tissue area of heart explants, we used Segment Anything for Microscopy (*77*) with the base model for natural images and user prompts to segment the explants. The segmented images were processed with a CellProfiler (*74*) pipeline that incorporated IdentifyPrimaryObjects and MeasureObjectSizeShape.

For investigating the effect of HAase on cardiac development *in situ*, eggs were opened after 48h of incubation. Embryos were then transferred to a culture dish using the filter paper method (*78*). Subsequently cardiac jellies were injected (see above) with either PBS or PBS with 2mg/ml HAase and transferred to the microscope (Cell Discoverer 37C, humid environment). Images of the heart region were taken every hour for a total of 22 hours. To investigate the effect of HAase on cardiac jelly, eggs were incubated for 26-29 (HH8-9) hours and transferred to a culture dish using the filter paper method. Coelomic cavities were injected (see above) with either PBS or PBS with 2mg/ml HAase and subsequently culture in a humidified incubator at 37C. After 18 hours of incubation, embryos were fixed and cryosectioned (see section below). *Cardia bifida* was induced by incision of the foregut after around 28 hours of incubation using micro scissors.

## Statistical analysis

Genotype comparisons (WT vs. HAS2 KO) were analyzed using unpaired t-tests that considered the mean value of each cell line. Hyaluronidase treatment window comparisons (untreated, d1.5-5.5, d2.5-5.5) were analyzed using paired t-tests with pairing performed across cell lines. For each cell line and treatment window combination, the mean value was calculated and used as the input for the paired t-test. P-values were adjusted for multiple comparisons using the Holm-Bonferroni FDR correction (α = 0.05). Timepoint comparisons (t_0_, t_1_, t_2_) were analyzed using paired t-tests with samples paired across timepoints to account for repeated measures from the same cardioid. Data from biological replicates (WT1, WT2, WT3) were pooled and used as the input for the paired t-tests. P-values were adjusted for multiple comparisons using the Bonferroni FDR correction (α = 0.05). Timepoint comparisons of heart explants were analyzed using paired t-tests with samples paired across timepoints to account for repeated measures from the same explant.

## Notes

### Summary of Updates

addition of Hyaluronic Acid molecular weight measurements (Fig. S1C)

